# Topology of synaptic connectivity constrains neuronal stimulus representation, predicting two complementary coding strategies

**DOI:** 10.1101/2020.11.02.363929

**Authors:** Michael W. Reimann, Henri Riihimäki, Jason P. Smith, Jānis Lazovskis, Christoph Pokorny, Ran Levi

## Abstract

In motor-related brain regions, movement intention has been successfully decoded from in-vivo spike train by isolating a lower-dimension manifold that the high-dimensional spiking activity is constrained to. The mechanism enforcing this constraint remains unclear, although it has been hypothesized to be implemented by the connectivity of the sampled neurons. We test this idea and explore the interactions between local synaptic connectivity and its ability to encode information in a lower dimensional manifold through simulations of a detailed microcircuit model with realistic sources of noise. We confirm that even in isolation such a model can encode the identity of different stimuli in a lower-dimensional space. We then demonstrate that the reliability of the encoding depends on the connectivity between the sampled neurons by specifically sampling populations whose connectivity maximizes certain topological metrics. Finally, we developed an alternative method for determining stimulus identity from the activity of neurons by combining their spike trains with their recurrent connectivity. We found that this method performs better for sampled groups of neurons that perform worse under the classical approach, predicting the possibility of two separate encoding strategies in a single microcircuit.

## 2 Introduction

Advances in experimental techniques have allowed us to record the *in-vivo* activity of hundreds of neurons simultaneously. This has gone along with new processing techniques to extract information from such spike trains. Some are based on the manifold hypothesis (Gallego et al., 2017), which describes the high-dimensional spiking activity of large neuron populations as determined by a lower-dimensional space whose components may be aligned with behavioral (such as movement direction) or stimulus variables. The reconstruction of the underlying space can then improve the decoding of such variables.

The manifold hypothesis posits that activity is limited to the lower-dimensional space due to constraints imposed by the synaptic connectivity (Gallego et al., 2017), however, details of this relation remain unclear. One identified constraint is that neurons with a large number of synaptic connections tend to spike at the same time as the population at large, while fewer connections allow neurons to spike when others are silent. Beyond this result, information on how the structure of synaptic connectivity shapes the neural manifold are scarce. This is in large part due to a lack of data: While a large number of neurons can be simultaneously recorded from, it remains challenging to determine the microstructure of their synaptic connectivity at the same time.

Conversely, in a model of a neural circuit, we have complete knowledge about connectivity as well as activity and can try to understand their relation. However, in order to study how connectivity shapes the structure of the neural manifold in biology, it is crucial that the connectivity of the model matches biology as closely as possible. Furthermore, to study this specific question, one additional requirement must be fulfilled: Encoding a lower-dimensional space in high-dimensional spike trains implies that the spiking activity is highly redundant. Such redundancy is often a means to overcome the presence of noise in a system (Tkacik et al., 2010). Indeed, there are various sources of noise affecting neural activity and it has been demonstrated to be often unreliable (Nolte et al., 2019; Tolhurst et al., 1983; Stern et al., 1997; Shadlen and Newsome, 1998). Consequently, a model to study the structural implementation of a neural manifold needs to include these noise sources, to fully understand its function.

One such model is the rat neocortical microcircuit model of Blue Brain (*NMC-model*, (Markram et al., 2015)). It includes detailed synaptic connectivity that replicates a number of biologically characterized features (Reimann et al., 2015; Gal et al., 2017) and recreates a diverse set of experimentally characterized features of cortical activity (Reyes-Puerta et al., 2015; Renart et al., 2010; Luczak et al., 2007). It models morphological detail in the form of 55 morphological types of neurons and type-specific synaptic transmission between them (Ramaswamy et al., 2015). Crucially, the model of synaptic transmission includes short-term plasticity (Stevens and Wang, 1995; Markram and Tsodyks, 1996; Abbott, 1997) and both spontaneous release (Simkus and Stricker, 2002; Ling and Benardo, 1999) and synaptic failure (Ribrault et al., 2011), both important sources of noise in neural microcircuit (Nolte et al., 2019; Faisal et al., 2008).

We first tested whether a lower-dimensional manifold could be used to describe the emergent activity of the model. To that end we used the model in a *classification task*: We subjected the model to a number of repeated synaptic stimulus patterns, recording the responses of all neurons. Then we used established techniques to reconstruct the neural manifold from the spiking activity and tested whether the values of its components were sufficient to classify the identity of the stimulus pattern injected at any given time.

Next, we investigated the dependence of classification accuracy on the connectivity of a neuronal population, thereby determining which local connectivity features give rise to a robust manifold capable of encoding information about the structure of a stimulus. We found significantly different numbers of motifs of three neurons in high-performing than in low-performing populations. To expand on this, we sampled neuronal sub-populations with uncommonly structured synaptic connectivity between themselves or to the rest of the population. Based on the results, a method to predict the classification accuracy of any sub-population based on a number of topological parameters was used. Finally, we developed an alternative method to reduce the dimensionality of spike trains that allowed us to reliably decode stimulus identity from the activity of populations where the classical technique failed.

## 3 Results

### 3.1 Detailed microcircuit model encodes stimuli in a lower-dimensional space

We began by investigating whether the manifold hypothesis, i.e. the hypothesis that spiking activity of a neuron depends on a linear combination of latent variables, can describe the results of simulations of the NMC-model. To that end we ran a simulation campaign subjecting 31,000 neurons in the model to eight different synaptic input patterns (Fig. 1A, B). The stimuli were injected by activating thalamo-cortical afferent fibers that in turn activated synapses placed onto modeled dendrites according to a layer-specific density profile from the literature (Meyer et al., 2010). Each stimulus activated a random 10% subset of the fibers with an adapting, stochastic spiking process for 200 ms (see Methods). This was immediately followed by the next stimulus, randomly picked from the eight patterns, resulting in an uninterrupted, random stream. Each repetition of a pattern used the same synaptic input fibers, but different randomly instantiated spike trains. In total, 4495 stimuli were presented, each pattern used 562 +− 4 (mean +− std) times.

**Figure 1:**
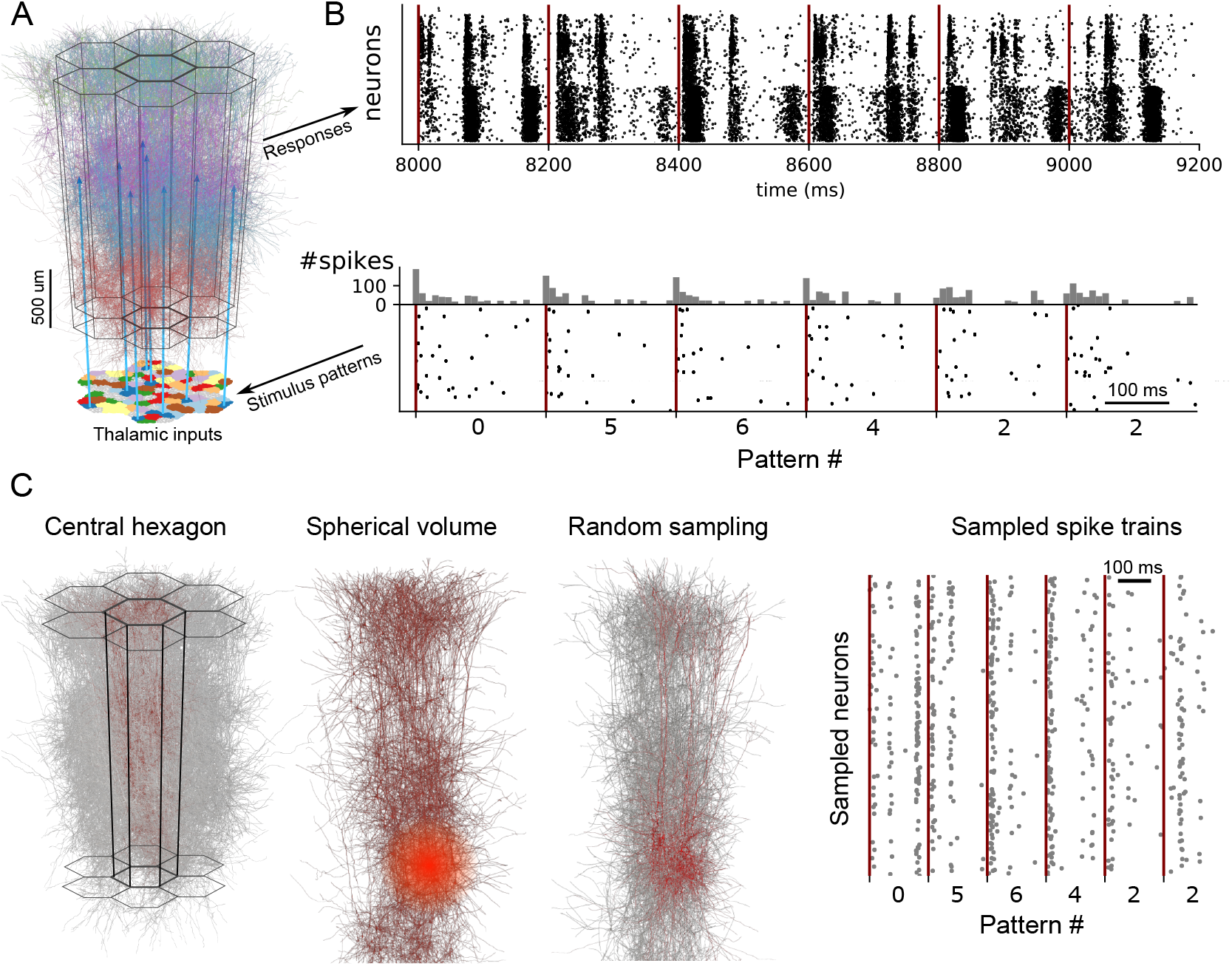
A: The modeled microcircuit includes 310 input fibers modelling thalamo-cortical afferents. B: Spike trains are prescribed to the fibers to serve as stimulus patterns. Each pattern is associated with 31 randomly selected fibers. Each repetition of a pattern, activates the associated fibers with an adapting Markov process, i.e. repetitions of a pattern use the same fibers, but with different stochastic spike trains. We apply a stream of 4495 repetitions of one of 8 stimulus patterns in random order (bottom) and simulate the microcircuits response (top). C: Random sampling: We sample at random offsets by randomly picking 600 neurons from all neurons within a given radius and consider their spike trains for further analysis.

To approximate the recording of spike trains with extracellular electrodes, we sampled neurons within a given radius from random offsets within the model and recorded their spike train for further analysis (Fig. 1 C). Initially, we performed 25 such samplings within 175*μm*. The number of samplings was set to conform with our second, structural sampling method presented in the next section. There most of our sampling parameters display a pronounced change in their distributions around the value 25 (Levi et al., 2020).

We then analyzed the samples according to the manifold hypothesis by first extracting the hidden components through factor analysis (time bin size 10 ms, first 12 components extracted, see Methods). We then investigated whether the values of these components can be used to distinguish between the stimuli. Exploratory analysis revealed that within three of the strongest components different stimuli followed different average trajectories (Fig. 2A), although individual trajectories were subject to a large amount of noise (Suppl. Fig. 5).

**Figure 2:**
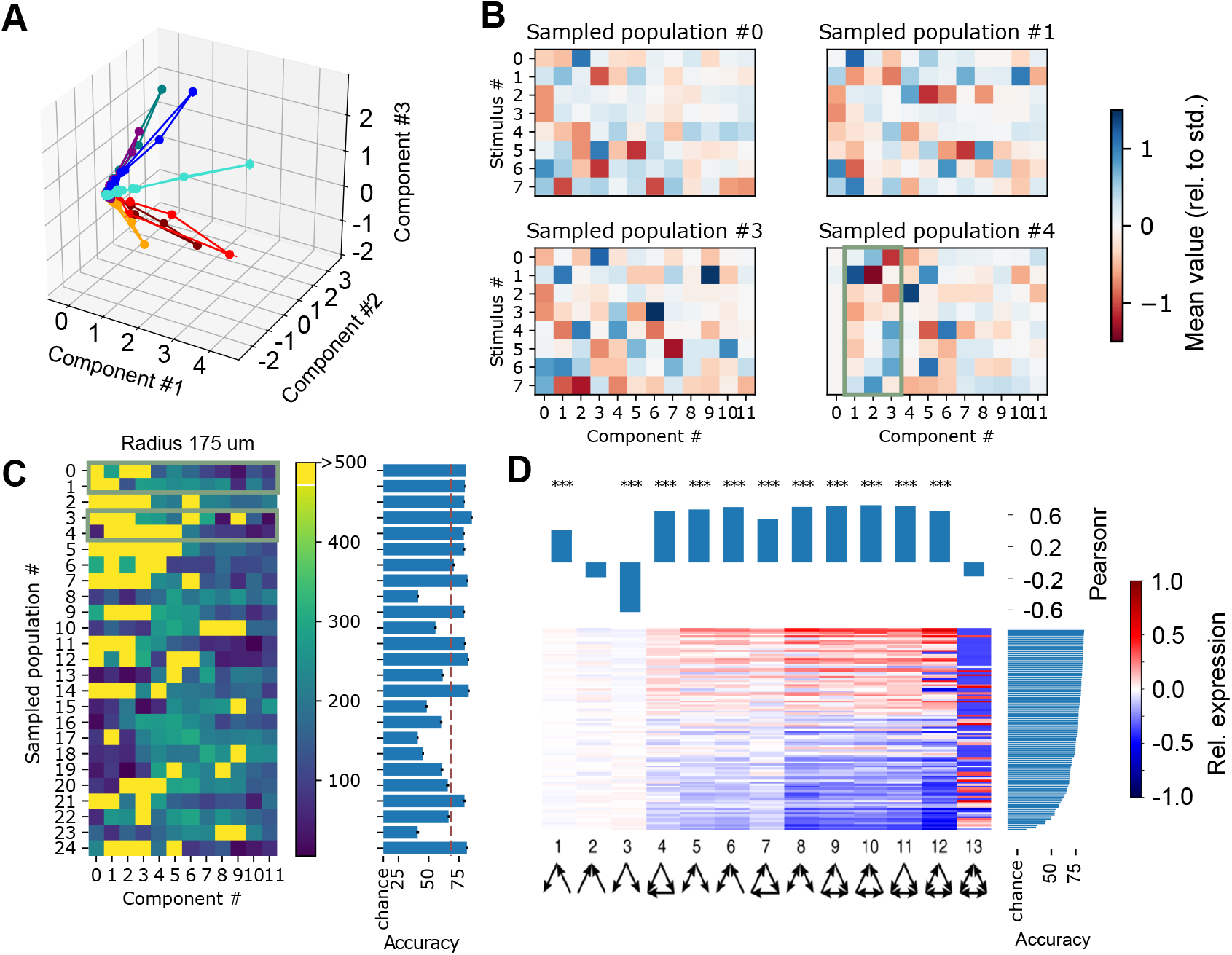
A: Trajectories of three components for an exemplary volumetric sample with a radius of 175*μm*. Responses to different stimuli are indicated in different colors (mean of 562 +− 4 (mean +− std) repetitions each). B: Mean of the most extreme value taken by individual components during a stimulus presentation, normalized with respect to the overall mean/std. Indicated for four random neuron populations, volumetrically sampled with a radius of 175*μm*. Green outline indicates the three components shown in A. C: (left) For 25 volumetric samples with radius 175*μm*: Negative logarithm of the p-values of a test against the null hypothesis that their extreme values as in B have identical means (Kruskal test). Green outlines indicate the samples shown in B. (Right) Mean accuracy of a classifier of stimulus identity based on the trajectories of the 12 main components of the indicated volumetric samples (see Methods). D: Bottom: triad motif over- and underexpression in 125 volumetric samples with radii between 125 and 325 *μm*, relative to ER. Samples are sorted by classification accuracy (right). Top: Correlation between expression of a specific motif and classification accuracy in the samples (pearsonr).

Next, we wanted to understand which of the twelve extracted components contained information about the identity of the presented stimulus. The time course of a component during a single stimulus followed a simple trajectory of rapidly rising to a maximum (or negative minimum) value and decaying back to zero (Fig. 2A). This allowed us to collapse the time course to the value furthest away from zero with minimal loss of information. We found that most of the time, this value differed significantly for the different stimulus patterns, with mean values for a given stimulus up to 1.6 standard deviations from the mean over all repetitions of all stimuli (Fig. 2B). In other cases however, components never strayed more than half a standard deviation away from the overall mean.

We tried to quantify these differences by calculating for each component its expected usefulness for distinguishing between stimuli. To that end, we calculated the negative logarithm of the p-value of a Kruskal test against the null hypothesis that the mean value was identical for different stimuli (Fig. 2C). This confirmed that pattern identity strongly affected the values for over half of the components in most volumetric samples. For some samples though, pattern identity affected only few and weaker components (Fig. 2C, samples #8, #23). Finally, we performed a classification task attempting to discern stimulus pattern identity from the trajectories of the first 12 components. The linear classifiers were 6-times cross validated with a split into 60% training data and 40% validation data (see Methods). We found that the resulting classification accuracies reflected the differences between the samples. Where stimulus identity significantly affected the trajectory of most components, accuracies exceeded 80%, where it only affected few or weaker components it could go as low as 40%. Yet, in all cases accuracies were significantly above the chance level of 12.5%.

We conclude that our simulations, combined with a volumetric sampling method lead to results that are comparable to the literature in the sense that different stimuli lead to different trajectories in a lower-dimensional manifold. However, the degree to which the manifold encodes stimulus identity varies between samples. According to the manifold hypothesis, the nature of the manifold is constrained by and therefore depends on the connectivity of the participating neurons (Gallego et al., 2017). We therefore aim to identify what feature of connectivity leads to more or less successful stimulus encoding.

We began by considering the presence of triad motifs in the volumetric samples. For each sample, we counted the occurrences of each possible motif of three connected neurons, and calculated their over- or under-expression compared to an Erdos-Renyi control model with the same number of neurons and connections (Figure 2D, bottom). When we compared the degree of expression of individual motifs to the classification accuracy, we found significant correlations for 11 out of 13 motifs. Specifically, for highly connected motifs with four or more connections, over-expression was associated with higher accuracy and under-expression with lower accuracy. For weakly connected motifs with only two connections, this trend was largely reversed. As the degree of expression of motifs was normalized to the total number of connections in the sample, this indicates that strongly clustered connectivity leads to increased accuracy in the classification task. Unfortunately, the observed trend was very similar for all highly connected motifs, ruling out more fine-grained observations. Therefore, we explored a different approach to study the effect of connectivity.

### 3.2 A classification task in samples with uncommon connectivity

The hypothesis that differences of the connectivity between volumetric samples explain their different accuracies in the classification task leads to the following prediction: A neuron sample with an uncommon pattern of connectivity is more likely to depict an outlying classification accuracy - either very low or very high. To confirm this, we used *chief-tribe* sampling to generate such samples with uncommon connectivity (Levi et al., 2020).

Briefly, any of the 31,000 neurons in the model can be considered a *chief* and its associated *tribe* contains in addition all neurons that are directly synaptically connected to it (Fig. 3A). To find tribes with uncommon connectivity, we defined 18 topological parameters that measure the structure of connectivity within, to, and from a tribe (see Table 1). We then sample the 25 tribes containing at least 50 neurons and with the highest values for a parameter as the *champions* of that parameter.

**Figure 3:**
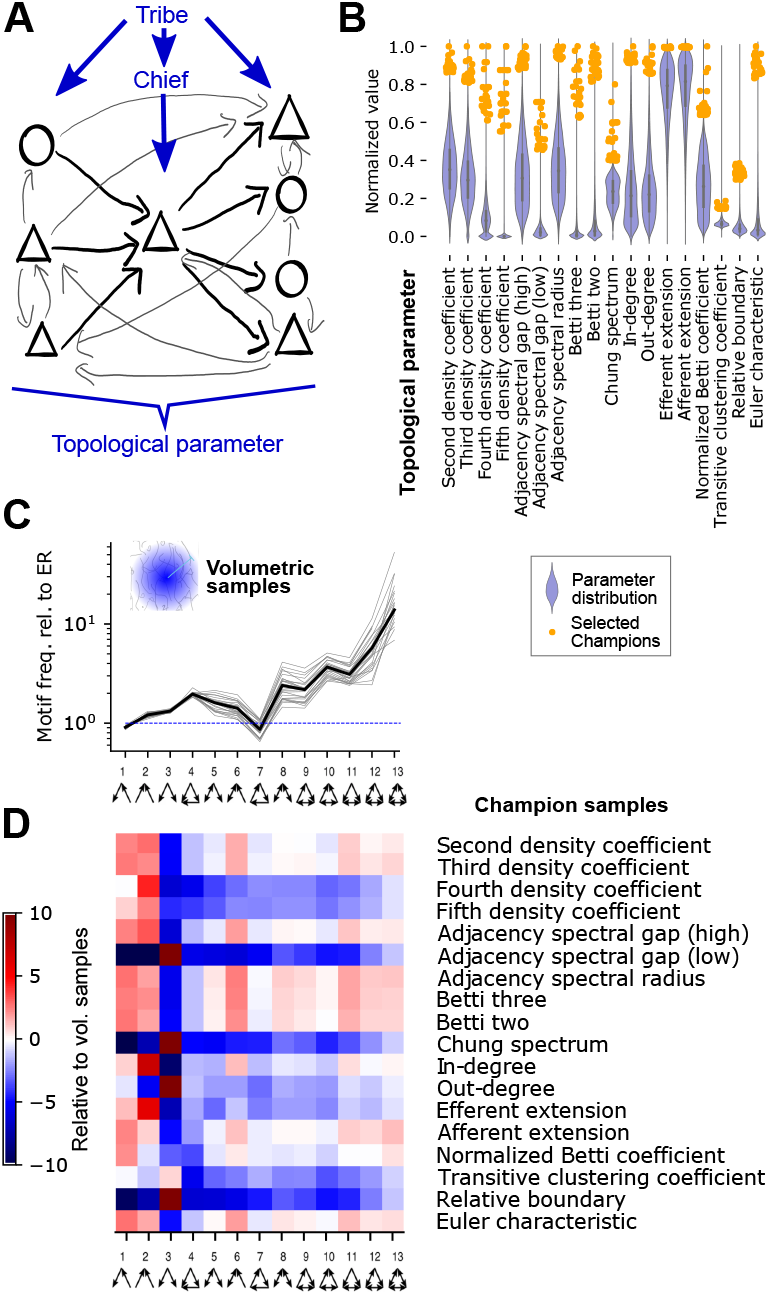
A: Chief-tribe sampling: Any neuron can be a *chief*. Its associated *tribe* are all synaptically connected neurons, plus the chief itself. We then calculate various topological parameters of the tribe’s connectivity. B: Blue violinplot: Distribution of various topological parameters, normalized between 0 and 1. Orange dots: Location of the *champions*, i.e. the 25 tribes of at least 50 neurons with the highest values for a parameter. C: Frequency of triad motifs in volumetric samples with a 175*μm* radius relative to an Erdos-Renyi graph with the same number of nodes and edges. Grey: 25 volumetric samples; Black: their mean. D: Over- and under-expression of triad motifs in the connectivity of *champion* samples of various parameters, normalized with respect to mean/std of the volumetric samples.

**Table 1:** Topological parameters used. Type denotes which connections determine the parameter: Only recurrent connections within the tribe, or additionally afferent connections into the tribe, or efferent connections. Parameters without a shorthand assigned were removed for being redundant. For details, see Methods.

| Parameter | Shorthand | Explanation | Type |
| --- | --- | --- | --- |
| Adjacency spectral gap (high) |  | Diff. between two largest moduli of eigenvalues of adjacency matrix | Recurrent |
| Adjacency spectral gap (low) | ASG | Lowest nonzero modulus of an eigenvalue of adjacency matrix | Recurrent |
| Adjacency spectral radius |  | Maximum modulus of an eigenvalue of adjacency matrix | Recurrent |
| Extension |  | Num. of neurons connected to but not in tribe |  |
| Afferent extension | AE | * for afferent connectivity | Afferent |
| Efferent extension | EE | * for efferent connectivity | Efferent |
| Betti numbers |  | Betti numbers of the tribes |  |
| Betti two |  | Second Betti number | Recurrent |
| Betti three |  | Third Betti number | Recurrent |
| Chung spectral gap |  | Smallest eigenval. of largest strongly connected component of tribe | Recurrent |
| Euler characteristic | EC | Euler charac. of the tribe | Recurrent |
| Density coefficients |  | Num. of k-simplices divided by num., of k-1-simplices and normalized |  |
| Second density coefficient |  | * for second dimension | Recurrent |
| Third density coefficient |  | * for third dimension | Recurrent |
| Fourth density coefficient | 4DC | * for fourth dimension | Recurrent |
| Fifth density coefficient | 5DC | * for fifth dimension | Recurrent |
| In-degree | ID | Total in-degree of the chief | Afferent |
| Out-degree | OD | Total out-degree of the chief | Efferent |
| Normalized Betti coefficient | NBC | Sum of normalized Betti numbers, | Recurrent |
| Relative boundary | RB | Number of connections into tribe, divided by connections within tribe | Afferent |
| Transitive clustering coefficient | TCC | Number of directed 3-cliques containing the chief divided by number of possible such cliques | Recurrent |

The champions of a parameter were indeed outliers, with values that fell far outside the bulk of the distribution of the parameter over all tribes (Fig. 3B). To confirm that the connectivity of the champion samples deviated from the overall structure of the model, we first quantified the over- and underexpression of triad motifs in the volumetric samples (Fig. 3C). We then compared the result to the expression of triad motifs in the champion samples and found large differences for all champions (Fig. 3D). For each champion the prevalence of at least one motif was over five standard deviations from the mean of the volumetric samples.

However, the profiles of over- and underexpression were very similar for some pairs of parameters, indicating that their associated champions were redundant. We therefore removed champions and parameters with too strongly correlating triad profiles, reducing the number of parameters to 11 (see Methods, Fig. 4, Table 1).

**Figure 4:**
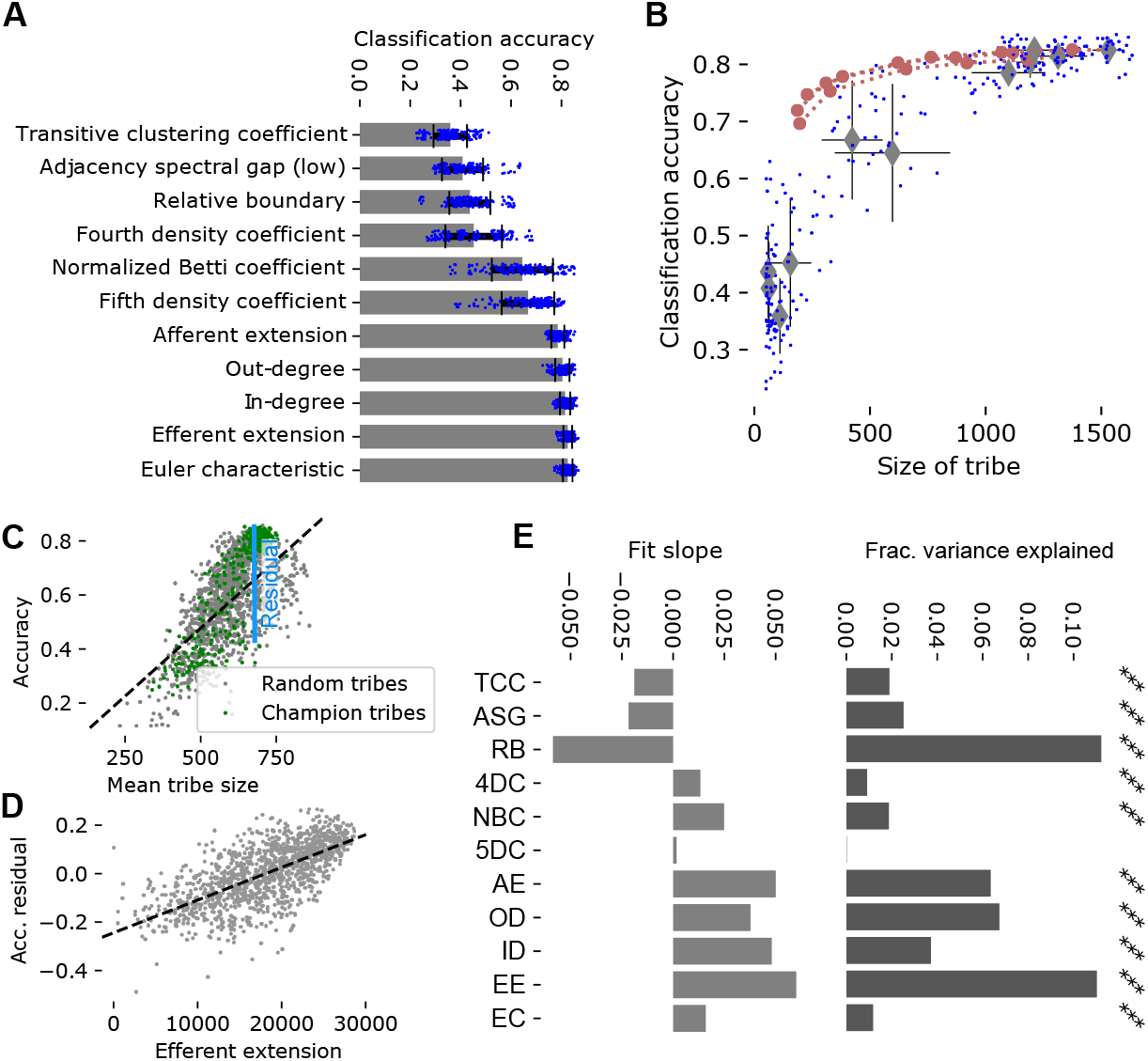
A: Accuracy of a linear classifier to decode stimulus identity from the spike trains of the champion-samples. Blue dots: Accuracies of 6-times cross validation for 25 champions (see Methods); bar and errorbars: mean and standard deviation. B: Size of champion tribes against their decoding accuracy. Blue dots: individual tribes; grey diamonds and errorbars: mean and standard deviation of champions of a parameter. Red circles: For the three largest classes of champions, subsampled at 90%, 70%, 50%, 25% and 15%. C: Mean size of associated tribes over neurons in champion (green) and randomly picked (grey) tribes. Black line: Linear fit; indicated in blue: residual of the fit. D: Efferent extension against the residual accuracy indicated in C for randomly picked tribes. Grey dots: 1276 random tribes; black line: linear fit. E: Left: Slopes of linear fits of residual accuracy against normalized values of the indicated parameter for randomly picked tribes. Right: Fraction of variance of the residual accuracy that is explained by the linear models over a model that takes only the morphological type of the chief into account (see Methods). Black stars indicate Bonferoni-corrected significance (***: *p <* 0.0001).

We then repeated the stimulus pattern classification task, but this time using spike trains from the champion samples. As predicted, the resulting classification accuracy varied drastically between champions of different parameters, possibly reflecting their differently structured connectivity (Fig. 4A). However, while the volumetric neuron samples always contained 600 neurons, the size of the champion samples depended on the connectivity of their chiefs and consequently varied. When we compared the size of the samples to the classification accuracies, we found a strong correlation (Fig. 4B, grey diamonds and blue dots).

This led to two questions: First, is this observed difference in accuracy a result of different connectivity – in this case, a lower combined in- and out-degree of the chief – or simply an artifact of the analysis – i.e. a consequence of using fewer spike trains in the classification task? And second, do the various topological metrics capture relevant features of connectivity beyond simply the size of the sampled tribe?

To address the first question, we repeated the pattern classification task for the three largest champions at reduced sizes. That is, we used only 90%, 70%, 50%, 25% or 15% of the neurons in the champion tribes of Euler characteristic, In-degree and Out-degree for classification (see Methods). The classification accuracies of these subsampled champions was only slightly reduced compared to the performance of the complete sample (Fig. 4B, red circles). Importantly, the subsampled champions performed at the top end or better than complete champions of comparable size. This indicates that for complete champions of smaller size (such as relative boundary), the lower number of spike trains being analyzed does not fully explain their reduced classification performance. Instead, the smaller size indicates weaker connectivity of the chief to the rest of the population, which may reduce the tribes ability to successfully encode stimulus identity.

To address the second question, we first generated a large number of additional tribal samples. This time, we sampled 1276 tribes by picking up to 25 chiefs randomly from each of the 55 morphological types of neurons in the model (Markram et al., 2015; Petilla Interneuron Nomenclature Group et al., 2008). We then subjected the samples to the stimulus pattern identification task (Fig. S6). This provided us with a large number of additional data points that were not biased towards extreme connectivity patterns, but represented typical connectivity of the model, while still providing variable classification accuracy. As before, classification accuracy in these samples depended on the sizes of tribes, in particular we found a linear relation to the *mean tribe size*, i.e. the mean over all neurons in a sample of the size of the associated tribe (Fig. 4C, grey dots). Corresponding data points for champion samples matched the fit, indicating that it captured the effect of tribe size we found earlier (Fig. 4C, green dots). We performed a linear fit of the data (Fig. 4C, black line) that allows us to predict the effect of tribe sizes on classification accuracy. Subtracting the prediction from the measured accuracies yields the *residual accuracy*, i.e. the accuracy with the effect of tribe sizes largely removed (Fig. 4C, blue). As the mean tribe size measure does not depend on the presence of a chief in the sample, we will later be able to generalize this concept to the volumetric samples.

We studied the relation between values of the various topological parameters and the residual accuracy using data from the randomly sampled tribes (for an example see Fig. 4D). Specifically, we compared the effect of only the morphological type of the chief to linear models combining the effects of morphological type and a topological parameter, calculating the additional fraction of variance explained by the combined model (Fig. 4E, right). This additional step let us rule out explanations where connectivity patterns captured by the topological parameters are merely correlated with the morphological types and have no direct effect on residual accuracy. Additionally we considered the slope of the linear fit in each parameter to assess the direction and strength of its effect. We found statistically significant effects for all parameters except the fifth density coefficient (t-test, *p <* 0.0001 Bonferroni-corrected; Fig. 4E). This indicated that the topological parameters were capable of measuring features of connectivity conducive to encoding the identity of a stimulus pattern in a lower-dimensional space.

### 3.3 Generalization for volumetric samples

Finally, we tried to generalize our ability to predict the classification accuracy from the topology of synaptic connectivity to the volumetric samples. As before, we calculated the mean of the sizes of tribes associated with neurons contained in the samples, but this time for volumetric samples with radii between 125*μm* and 325*μm*. Based on this, we predicted their classification accuracies using the linear fit from before, and subtracted the prediction to gain residual accuracies. We then calculated the values of the topological parameters based on the connectivity of the samples (Fig. 5A, left) and analyzed their relation to the residual accuracies as in the previous section. However, the parameters in-degree, out-degree and transitive clustering coefficient were defined with respect to the chief of a tribe (see Table 1) and consequently undefined for volumetric samples. We instead used the mean of in-degree, out-degree and clustering coefficient over all neurons in a sample. Most parameters calculated this way had no significant effect on the residual accuracies with only three exceptions (Fig. 5B). It thus appears as if the ability of most topological parameters to capture significant connectivity patterns is linked to tribal structure of the samples we used before.

**Figure 5:**
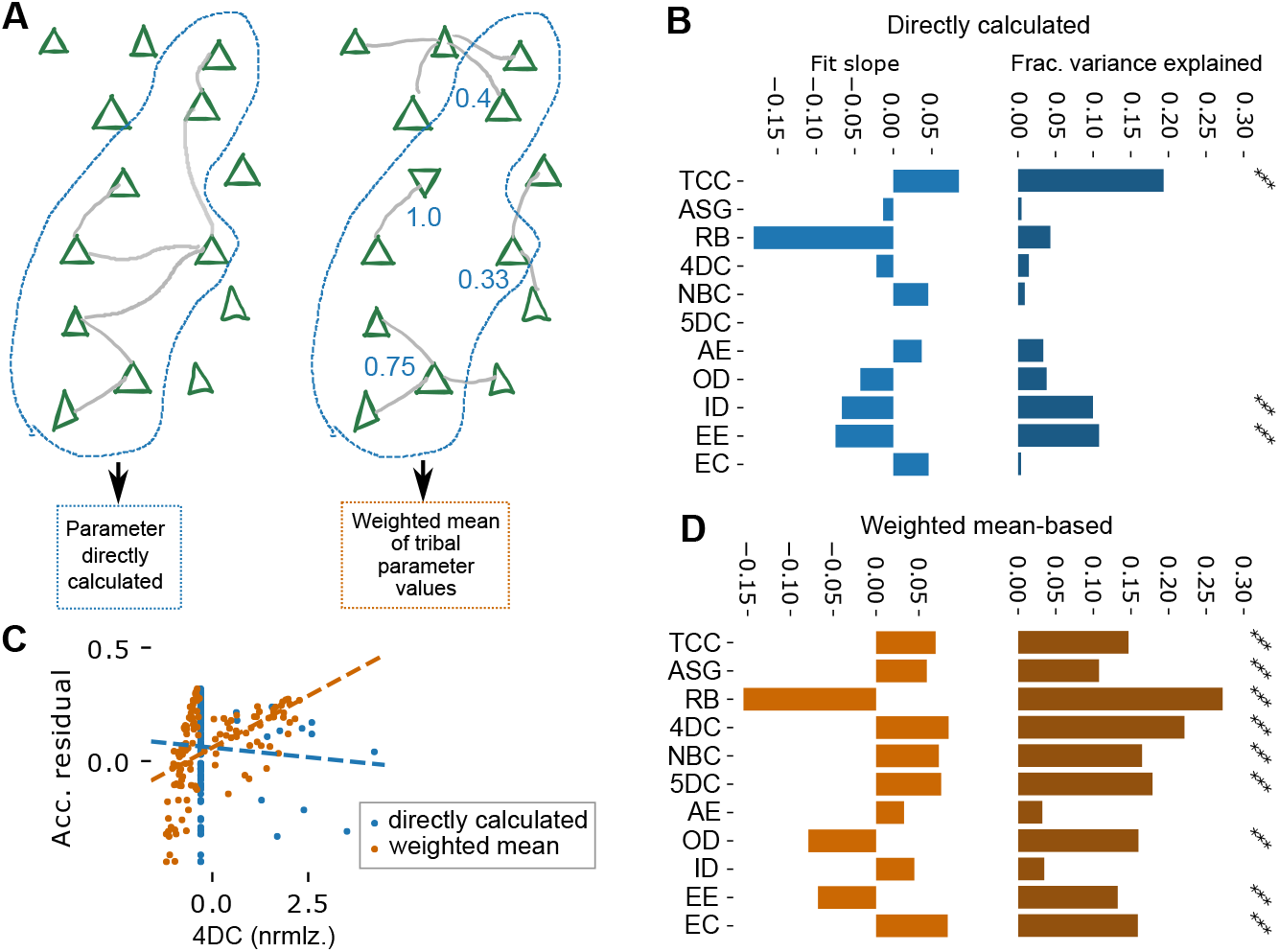
A: Two ways of calculating values for topological parameters in volumetric samples. Left: Values are calculated directly on the synaptic connectivity within the sample. Right: Generation of “synthetic” values of topological parameters that take the tribal structure of the sample into account: The size of the overlap of each possible tribe with the sample is calculated, then a weighted average of the parameter value for the *N* strongest overlapping tribes is used. B: Left: Slopes of linear fits of residual accuracy against normalized parameter values that were directly calculated. Right: Fraction of variance of the residual accuracy that is explained by the linear models over a model that takes only the radius of the sample into account (see Methods). Black stars indicate Bonferoni-corrected significance (***: *p <* 0.0001). Shorthand parameter names see Table 1. C: Comparing directly calculated parameter values (blue) to weighted mean-based, “synthetic” values of the fourth density coefficient. D: As in B, but for synthetic values of topological parameters.

Therefore, we developed the following method to generate a *synthetic value* of a topological parameter for a volumetric sample that takes its tribal structure into account (Fig. 5A, right). First, calculate the relative size of the overlap between each tribe and the sample. Next, find the *N* tribes with the strongest overlap. Finally calculate the synthetic value as the weighted mean of parameter values of those *N* tribes, where the weights are proportional to the relative overlap sizes. For each topological parameter, an optimal value for *N* was determined that maximized the correlation between the resulting synthetic values for volumetric samples and their residual classification accuracies (see Fig. 5C for an exemplary result). Using this method, all but two investigated parameters provided a statistically significant effect (t-test; *p <* 0.0001 Bonferroni-corrected; Fig. 5D; Fig. S8A), reinforcing the idea that differences in classification accuracy can be explained by the connectivity of the sampled neurons, and the topological parameters capture some of the features of connectivity that are relevant for this.

The optimal number of tribes to include in the weighted mean (*N* above) tended to be low, under 300 tribes for most parameters, with larger numbers leading to a gradual decline in correlation with classification accuracy (Fig. S8B). Exceptions were the *transitive clustering coefficient*, *efferent extension* and *out-degree*, where the optimal solutions were weighted means including all 31,000 tribes. Curiously, this was accompanied by a switch of the sign of the correlation as more tribes were included (compare Fig. 4E to Fig. 5D). This indicates that at least these two classes of parameters capture different ways in which connectivity affects the performance of a neuron sample in the classification task.

### 3.4 The role of low-performing populations

Having identified the connectivity patterns that improve a population’s ability to encode stimulus patterns, the question remains what the role of the low-performing populations might be in the neural computations of a microcircuit. These are populations containing neurons forming small tribes, that is, neurons with structurally weak coupling to the rest of the population. A dichotomy between weakly and strongly coupled neurons has been characterized before in the form of *soloists* that spike when the rest of the population is silent, and *choristers* that spike together with the rest (Okun et al., 2015). We therefore investigated whether the low-performing samples and tribes contained a larger amount of soloists.

We began by calculating the *coupling coefficient* of all excitatory neurons in the population and comparing its distribution to two controls with shuffled spike trains: One that preserved only the overall firing rate of the entire population and one that preserved firing rates of individual neurons (see Methods, Fig. 6A). We confirmed that in our simulation both soloists and choristers, that is, neurons with an unexpectedly low or high coupling coefficient emerged. Next, we investigated whether this property depended on the size of the tribe that a neuron is the chief of (Fig. 6B). We found a strong, positive correlation between these properties, albeit only for chiefs in layers 1 to 5. Curiously, for chiefs in layer 6 the relation reversed, with larger tribes apparently reducing the coupling coefficient of their tribe. At the same time, neurons in layer 6 depicted the highest coupling coefficients.

**Figure 6:**
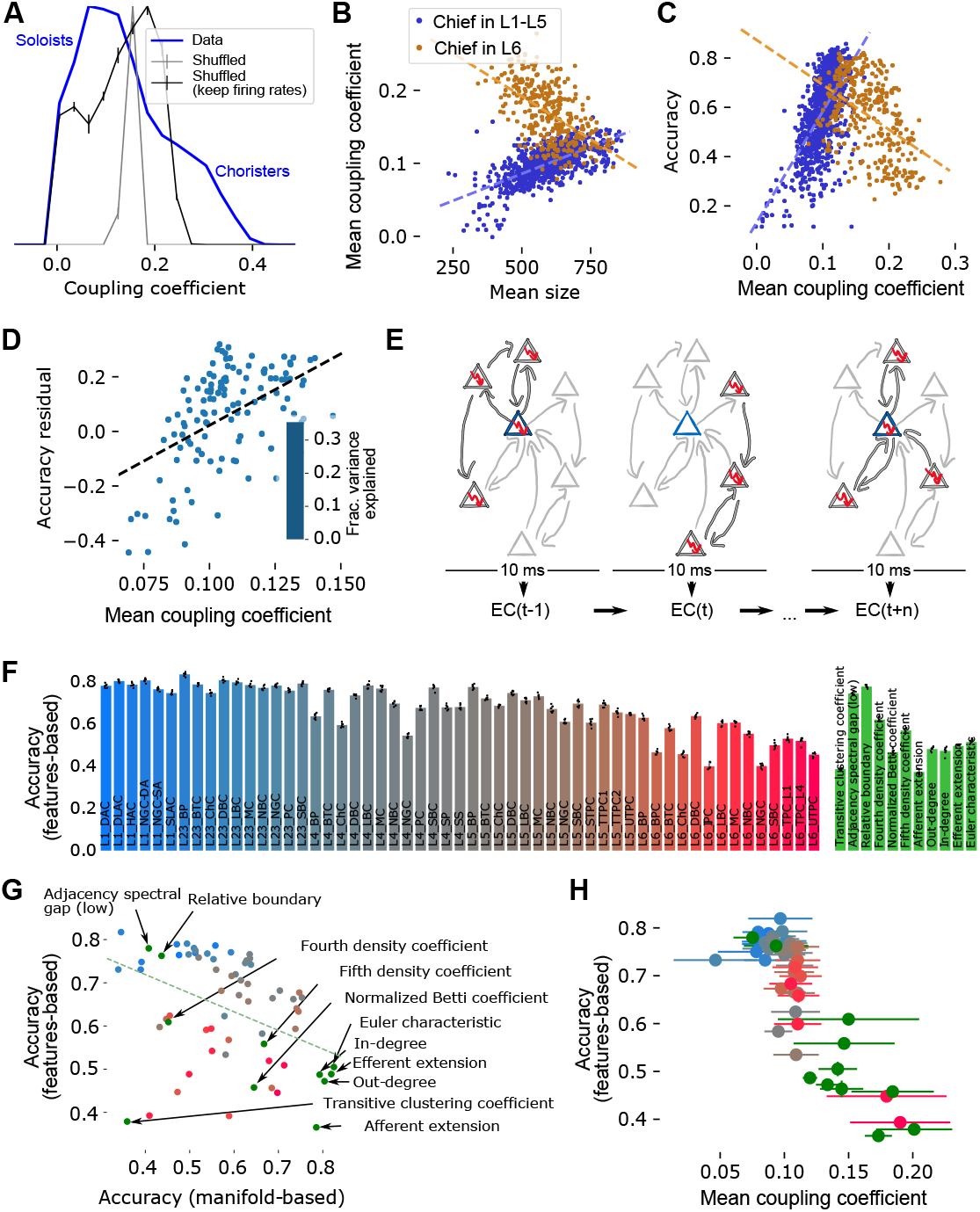
A: Distribution of the *coupling coefficient*, as in Okun et al., 2015, of all excitatory neurons in the model. Blue: data; grey lines with errorbars: Shuffled controls that keep or shuffle mean firing rates of individual neurons. Neurons with an unexpected low coefficient are “soloists”, a high coefficient indicates a “chorister”. B: Tribe size against the coupling coefficient of the chief for chief neurons in layers 1-5 (blue) and in layer 6 (orange). Dashed lines: linear fits. C: As in B, but coupling coefficient against classification accuracy of the tribes. D: Mean coupling coefficient of neurons in a volumetric sample against its residual classification accuracy, as in Fig. 5C. Inset: Fraction of variance explained as in Fig. 5D. E: Classification based on *topological featurization*: For a given tribe, we consider in each 10 ms time step only the subset of neurons that are firing. We then calculate the value of the Euler characteristic (EC) of the graph of connectivity between them. We classify stimulus identity based on the EC time series of 25 chiefs. F: Left: Results of the classification for chiefs randomly picked from the various morphological types in the model. Errorbar: std of cross validation. Right: Same, for the various champions. G: Mean classification accuracies using the manifold-based method against accuracies based on topological featurization. Red/blue dots: Random chiefs as in F; green dots as labelled. H: Accuracies as in F against the mean coupling coefficient of neurons in the tribes. Errorbars: std over coupling coefficients.

Having found that the size of a tribe affects both classification accuracy and coupling coefficient, it was no surprise that we also found a strong correlation between the coupling coefficients of neurons in a tribe and its classification accuracy (Fig. 6C). However, once again, tribes with chiefs in layer 6 reversed this correlation. For the volumetric samples, we analyzed the effect of mean coupling coefficient on residual accuracy, finding the same strongly positive correlation as for layers 1 to 5 (Fig. 6D). The coupling coefficient explained over 30% of the variance of residual accuracy, more than any of the topological parameters (compare inset to Fig. 5D). Taken together, this confirmed that neurons with an uncommonly low coupling coefficient below 0.1 were more prevalent in tribes and volumetric samples that did not perform well in the classification task.

In order to further explore the ways in which stimuli are encoded in microcircuit activity, we employed an alternative decoding method based on *topological featurization* (Fig. 6E Levi et al. (2020)). Similar to the manifold-based method, it begins by binning the spike trains of individual neurons in a tribe into 10-ms time bins. However, the way the dimensionality of this result is reduced, differs drastically. In each time bin, we consider all neuron members of a tribe that are spiking, and the connections between them. We then calculate the Euler characteristic (*EC*) of the flag complex of the graph of the spiking sub-network (see Section 5). This yielded a time series of *EC* values that represent the time course of activity in the tribe. As this reduced the dimensionality to a single dimension with ten time steps for a tribe, we grouped 25 tribes together and used their pooled time series in the same linear classification method as before. The tribes pooled together were either champions of the same parameter, or their chiefs randomly picked from the same morphological type.

As for manifold-based classification, the results depended on the type of tribe. For pooled tribes based on picking chiefs randomly from a given morphological type, we found an overall gradient from accuracy around 80% for chiefs in superficial layers, to under 50% for chiefs in layer 6 (Fig. 6F, left). For pooled champions, the accuracy varied comparably between 40% and 80%, depending on the topological parameter they were champion of (Fig. 6F, right).

Astonishingly, the results were almost completely the inverse of the manifold-based classification: Tribes that performed well in the manifold performed poorly for topological featurization and vice versa (Fig. 6G). Only some tribes with chiefs in layer 6 and the champions of the transitive clustering coefficient performed equally badly for both methods. This indicated that topological featurization is a classification method more suitable for the samples that performed poorly in manifold-based classification, that is, samples with a large number of soloist-type neurons. Indeed, we found a strong negative correlation between the mean coupling coefficient of a tribe and its topological featurization-based classification accuracy (Fig. 6H). We therefore predict that soloists employ an alternative scheme for encoding information in their spike trains; one that can be more readily read out when in addition to their spiking activity their synaptic connectivity is taken into account.

While we have demonstrated that topological featurization performs good classification for soloists, the question remains: Why does it fail for choristers? One of the defining features of choristers was their overall larger degree, leading to large associated tribes. We therefore hypothesized that these larger tribes were also more strongly overlapping, leading to more similar *EC* time series. As such, the information in the time series would be more redundant – each tribe conveying the same information. To test this, we further investigated to what extent the 25 champion samples belonging to the same parameter were overlapping and how similar their *EC* time series were. To that end, we computed the mean relative overlap of pairs of tribes using the Szymkiewicz-Simpson coefficient (Vijaymeena and Kavitha, 2016), i.e. the relative number of common neurons shared by a pair of tribes. Likewise, we computed the mean correlation coefficients (Pearson) between all pairs of pooled *EC* time series averaged over trials, referred to as *feature correlation*. We found a strong positive correlation between feature correlation and samples overlap in champion samples (Fig. 7A), indicating that a higher number of shared neurons is linked to a higher correlation in the corresponding feature time series. Two examples of pairwise feature correlation matrices of the parameters with the lowest and highest mean correlations respectively are illustrated (Fig. 7B). Our next question was whether such a redundancy in the feature time series would lead to reduced classification accuracies. And indeed we found a negative correlation between featurization-based mean accuracies (over 6-times cross validation) and mean feature correlations (Fig. 7C).

**Figure 7:**
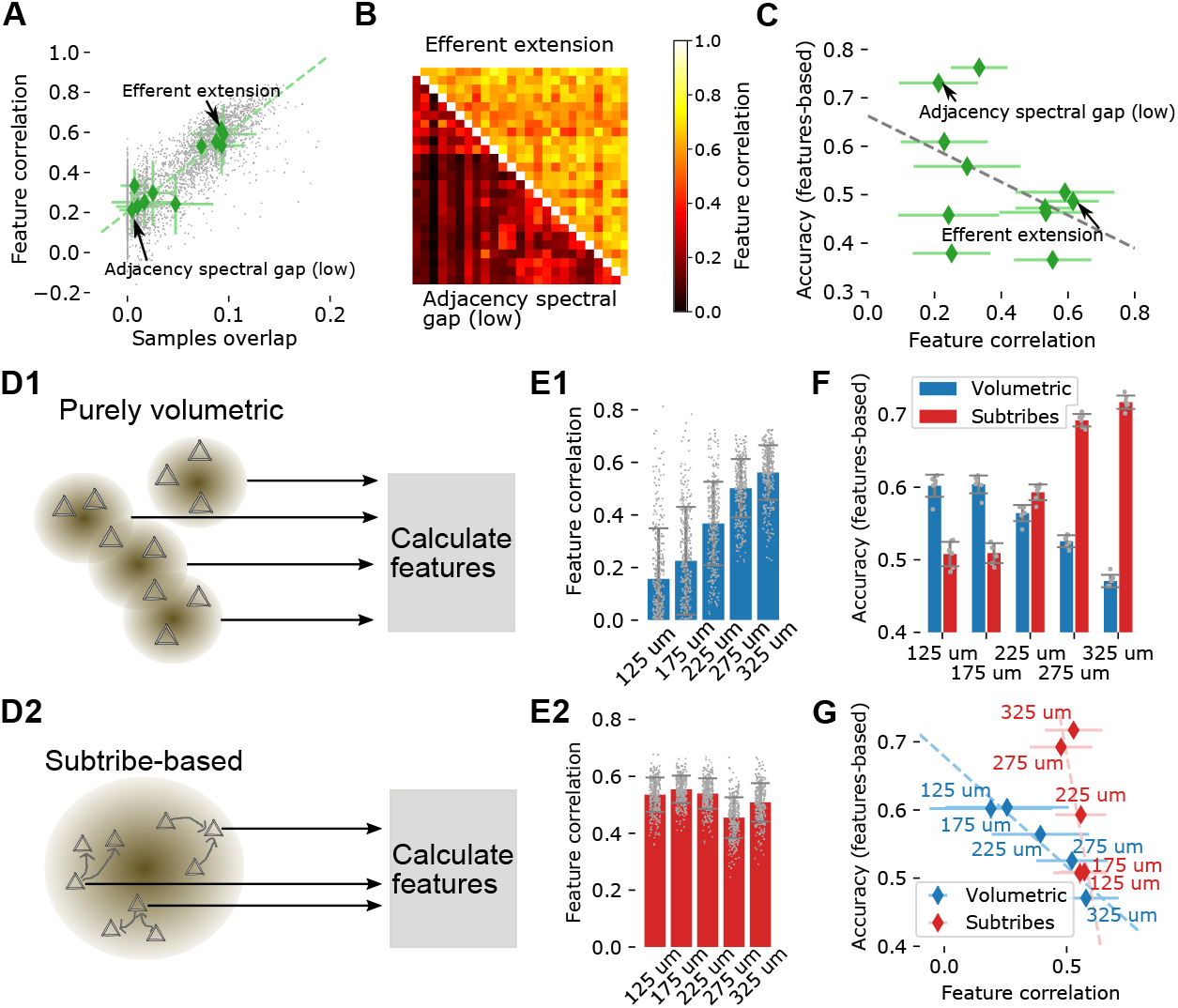
A: Mean relative overlap of pairs of tribes that are champions of the same parameter against the mean correlation coefficient of their Euler characteristic time series (*feature correlation*. B: Pairwise feature correlations of the champions of two exemplary parameters. C: Mean feature correlation of champions against their performance in feature-based classification. D1: Feature-based classification for volumetric samples can be performed using the active subnetworks of each sample in each time step, then pooling over samples. D2: Alternatively, the 25 largest tribes can be found within a single volumetric sample, which are then analyzed as other tribal samples. E1: Mean feature correlation for volumetric samples of various radii when the technique in D1 is employed. E2: Same, when the method outlined in D2 is employed. F: Feature-based classification accuracies for volumetric samples using the purely volumetric (blue) or subtribe-based (red) techniques. G: Feature correlation against classification accuracy for volumetric samples using the purely volumetric (blue) or subtribe-based (red) technique.

As before, our next aim was to apply this technique to the volumetric samples. A straightforward approach to this simply uses the active sub-networks of 25 samples, pooling their *EC* time series (Fig. 7D1). However, as we found earlier that taking the tribal structure of a sample into account led to better results, we also used an alternative approach: Using only the neurons in a single volumetric sample, we extracted the 25 largest tribes that could be found within a single volumetric sample, referred to as *subtribes*, which were then analyzed as the other tribal samples (Fig. 7D2).

In the first approach, when applying the featurization directly on volumetric samples, we found a clear dependence of the mean feature correlation on the sampling radius, with larger radii resulting in higher feature correlations (Fig. 7E1). This was expected, as larger radii naturally resulted in larger overlaps of samples volumes. In the second approach, when applying featurization on subtribes, the resulting feature correlation seemed to be rather independent of the sampling radius (Fig. 7E2).

Astonishingly, when comparing the featurization-based classification accuracies of both approaches, we found that the subtribes-based approach clearly outperformed the volumetric approach for large radii (Fig. 7F). We found opposite dependencies of the mean accuracies on the sampling radii, reaching highest values for large radii in the subtribes-based approach and for small radii in the volumetric approach respectively. So, although the subtribes-based approach only used information extracted from a single volumetric sample instead of pooling several of them, it clearly performed better above 225*μm*. As before, we again found a negative correlation between accuracies and feature correlation values for both approaches (Fig. 7G). Taken together, these results suggest that features are better predictors when taking the tribal structure within a volume into account than just combining features from multiple volumes.

## 4 Discussion

We have confirmed that the emergent activity of the NMC-model can be described according to the manifold hypothesis, that is, as determined by a lower-dimensional space; and that the values of those dimensions encode information about inputs given to the model. We have further demonstrated that within the morphological, electrical and synaptic diversity captured by the model, some neuronal populations are more useful in reconstructing the lower-dimensional space and using it to decode information about a stimulus. We found that these differences can be largely explained by the synaptic connectivity of neurons in the populations.

A large component of this finding was that neurons that are strongly connected to the rest of the population tend to be of the “chorister” type (Okun et al., 2015) and provide better capability for decoding stimulus identity, when using manifold-based techniques. This is not surprising, as a chorister is defined by spiking together with many other neurons and is consequently aligned with the stronger components of the underlying manifold. Thus, their spike trains carry information about those strong components.

However, we also found topological features of connectivity influencing the manifold that go beyond merely adding more synaptic connections and instead capture the specific structure of them. We demonstrated that topological parameters such as density coefficients and clustering coefficients can be used to predict the success of a stimulus classification task. The clustering coefficient measures the tendency of the connectivity to form tightly bound assemblies and the density coefficients measure the tendency to form specific types of clusters called directed simplices (Reimann et al., 2017). Their effect can be explained as follows: While synaptic input strongly constrains neuron activity, this effect is weakened by the presence of noise, especially synaptic noise. At the same time, it has been demonstrated that correlations in synaptic input diminish the effect of noise (Wang et al., 2010). This means that a neuron participating in connection motifs that generate correlated input, will be less subject to noise, and constrained more tightly to the manifold, improving the value of its spike train in decoding the manifold. Directed simplices have been shown to generate such correlations before (Nolte et al., 2019).

We found that the distinction between “well-classifying” and “poorly-classifying” neurons parallels the split into “soloists” and “choristers” that has been found before (Okun et al., 2015). When we further investigated the emergent activity, we employed an alternative, topology-based method to reduce the dimensionality of spike trains of large neuronal populations (Levi et al., 2020). With this “topological featurization” technique, previously poorly-classifying neurons provided sufficient information to classify stimuli with high accuracy. While this technique cannot currently be employed *in-vivo*, as it requires information about the local connectome, advances in connectome reconstructions from electron-microscopy will allow us to validate this method in the future.

Topological featurization required the pooling of information from multiple neuron samples, unlike the manifold-based approach that kept them separate. Therefore, it arguably used more information for classification, potentially explaining its superior performance under some circumstances. This is however contradicted by the fact that the samples performing well with featurization were the ones that were individually small, and the larger samples performed more poorly. Further, for volumetric samples, the approach that took only a single sample instead of pooling over 25 samples had superior performance. Finally, we found that neuron samples classifying well with the featurization approach tend to be “soloists”. Taken together, this indicates the presence of an alternative encoding scheme employed specifically by “soloist”-type neurons.

Does this mean that there exists neural circuitry that reads out the Euler characteristic of another population? While such a claim would certainly be premature, we at least want to consider whether such circuitry could be implemented with known biological mechanisms. The Euler characteristic is the sum of the counts of directed *n*-cliques for increasing *n* with alternating signs (see Methods). This would require, for each *n*-clique *σ* one additional readout neuron *b_σ_* receiving input from all neurons in *σ*. Where *b_σ_* should fire only when receiving input from all neurons in *σ* simultaneously, forming conceptually an AND conjunction and indicating that the entire *n*-clique is active. This could be achieved through clustering of inputs from *σ* onto adjacent locations on the dendrite of *b_σ_* such that their concurrent activation triggers dendritic nonlinearities while individual activations remain local. Both cooperativity between nearby synapses (Harnett et al., 2012; Weber et al., 2016) and functional clustering of synapses with similar receptive properties (Iacaruso et al., 2017) has been observed experimentally. Topologically, *b_σ_* would extend the *n*-clique to an *n* + 1-clique and indeed we have found before that *n*-cliques are part of an unexpectedly high number of *n* + 1-cliques (Reimann et al., 2017).

The final readout *EC* would be a neuron receiving input from all *b_σ_*s, with a sign that depends on the size (value of *n*) of the clique that it reads from (see Fig. S9). That would mean that readouts from even-sized cliques would all need to be inhibitory, readouts from uneven-sized cliques excitatory. There is no evidence for such strict motifs, nor for mechanisms that would lead to their formation. But it is possible that a less strictly alternating sum could provide similar information as the Euler characteristic. In summary, a neural readout circuitry implementing the Euler characteristic is possible, but only some of its possible building blocks have been found experimentally.

These results were obtained in a highly detailed model, albeit one of only an individual microcircuit acting alone. The nature of biological neural manifolds is likely determined as well by long-range connections and dynamic interactions between different brain regions, meaning a purely local view is limited (Allen et al., 2019; Stringer et al., 2019). Yet, our results provide predictions on the general principles that may be employed to encode information in lower-dimensional neural manifolds.

## 5 Methods

### 5.1 Network simulations

Simulation methods are based on a previously published model of a neocortical microcircuit of the somatosensory cortex of the two week-old rat, here called the *NMC-model* (Markram et al., 2015). Synaptic connectivity (with apposition-based connectome) between 219,422 neurons belonging to 55 different morphological types (m-types) was derived algorithmically starting from the appositions of dendrites and axons, and then taking into account further biological constraints such as number of synapses per connection and bouton densities (Reimann et al., 2015). Neuronal activity in the NMC-model was then simulated in the NEURON simulation environment (www.neuron.yale.edu/neuron/). Detailed information about the circuit, NEURON models and the seven connectomes of the different statistical instantiations of the NMC-model analyzed in this study are available at bbp.epfl.ch/nmc-portal/ (Ramaswamy et al., 2015). Simulations and analysis were performed on an HPE SGI 8600 supercomputer (BlueBrain5).

We simulated evoked activity of the model described above for 900 seconds. After one simulated second (at *t* = 0 ms, as we discard the first second) we start applying a thalamic stimuli through synapses of 310 VPM fibers that innervate the microcircuit. The stimuli are a stream of eight different input patterns, where each pattern activates 31 randomly picked VPM fibers for 75 ms with an adapting, stochastic spiking process starting at 75 Hz and decaying to zero with a time constant of 20 ms. The stream consisted of repetitions of the eight patterns in random order, presenting one pattern every 200 ms. Repetitions of the same stimulus used the same fibers, but with different stochastic instantiations of the spiking process. Additionally, each of the 310 fibers was activated with a poissonian firing rate of 0.2 Hz. As for each stimulus the pattern was chosen randomly, the total numbers of presentations differed slightly between patterns: Pattern 0: 558 repetitions; pattern 1: 559; pattern 2: 566; pattern 3: 568; pattern 4: 561; pattern 5: 560; pattern 6: 557; pattern 7; 566.

### 5.2 Topological methods

Abstractly, the brain can be viewed as a *directed graph* 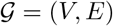, consisting of a set of *vertices V*, corresponding to the neurons, and a set of *directed edges E*. The elements of the edge set *E* are all pairs (*i, j*), where *i* and *j* are vertices in *V*, such that there is a synaptic connection from neuron indexed *i* to neuron indexed *j*. Reciprocal edges between any pair of vertices *i* and *j* (namely both pairs (*i, j*) and (*j, i*)), are allowed, but no more than one edge in the same direction. Loops, namely pairs (*i, i*) corresponding to edges whose start and end point is the same are also excluded.

If 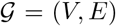 is a directed graph and *v ∈ V* is a vertex, then the *closed neighbourhood* of *v* is the set of all vertices in 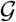 that are connected to *v* (in either direction) and the vertex *v* itself. The subgraph of 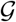 that consists of the vertices in the closed neighbourhood of *v* and all edges between them is also referred to in the mathematical literature as the *closed neighbourhood*, or the *closed neighbourhood graph* of *v*.

A collection of *n*-vertices *v*_1_*, …, v_n_* ∈ *V* of a graph in which every pair of vertices is connected by an edge is called an *n-clique*. If 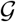 is a directed graph then a subgraph on *n* vertices such that every pair of them is connected by an edge in 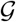, and which does not contain as a subgraph a 3-clique that is oriented cyclically, is called a *directed n-clique*. If 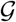 is a directed graph then one can construct a topological space that is made out of the directed *n*-cliques in 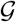. This topological space is called the *directed flag complex of* 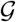 (Reimann et al., 2017). If *v* is a vertex in a directed graph 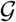, then both the closed neighbourhood graph of *v* and its associated directed flag complex are referred to as the *tribe of v*. For a more comprehensive mathematical explanation of these concepts and the associated invariants the reader is referred to Levi et al. (2020). We now describe the parameters from Table 1 in a topological setting. In the Supplementary Material 9.1 we give more heuristic explanations to complement the mathematical definitions.

### 5.3 In-degree and out-degree

If 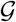 is a directed graph and *v* is a vertex in 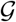, the *in-degree of v in* 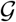 is the number of directed edges in 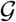 that end at *v*, and the *out-degree of v in* 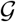 is the number of directed edges that emanate from *v* in 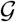. The *degree of v in* 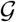 is the sum of its in-degree and out-degree. Note that the degree of *v* in 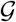 is one less than the number of vertices in the closed neighbourhood of *v*. Degrees of all the vertices in a graph, and associated degree sequences, characterize properties of the graph (Diestel, 2017). In- and out-degree of a single vertex in a graph are, however, very local parameters, and in themselves do not inform on global structure of the graph. Note that a directed *n*-clique is characterized by having one vertex with in-degree zero (within the clique), called the source, and one vertex with out-degree zero (within the clique), called the sink.

### 5.4 Transitive clustering coefficient

The *clustering coefficient* at a vertex *v* in a graph 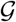 is a measure of how interconnected the vertices of the neighbourhood of *v* are. This coefficient has been widely studied, initially by Watts and Strogatz (Watts and Strogatz, 1998) in undirected graphs and by Fagiolo (Fagiolo, 2007) for directed graphs. We use a variation called the *transitive clustering coefficient*, which appeared in Levi et al. (2020), and is more suitable to our context, since in our topological analysis we only consider directed cliques in the construction of the directed flag complex, and therefore in tribes. The transitive clustering coefficient of a vertex *v* in a directed graph 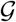 is defined by

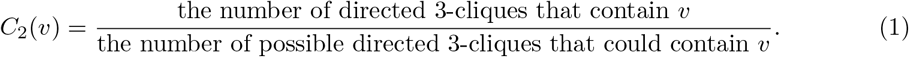

Any pair of edges that are incident to *v* could theoretically form one or two directed 3-cliques depending on their orientation. Therefore, the numerator in the definition is the number of actual directed 3-cliques that contain *v*, while the denominator is the maximum number of theoretically possible directed 3-cliques that contain *v*. The difference between this definition and that of Fagiolo is that in his definition all possible 3-cliques are considered (including the cyclical ones), where in our definition the cyclical cliques are omitted from the count. The number of possible directed 3-cliques at a vertex *v* in a directed graph is calculated as a function of the in-degree and out-degree of *v*, as well as the number of reciprocal connections it forms (Levi et al., 2020).

Topologically a 3-clique is 2-dimensional, hence the 2 in the subscript of *C*_2_. The transitive clustering coefficient can be generalised for higher dimensions, but this is not used here.

### 5.5 Density coefficient

Every directed (*k* + 1)-clique contains *k* + 1 directed *k*-cliques as its *faces*, but no number of directed *k*-cliques will necessarily form any (*k* + 1)-cliques. If we let *Q_k_*(*v*) denote the number of directed *k*-cliques that contain a vertex *v*, then the quotient *Q_k_*_+1_(*v*)*/Q_k_*(*v*) gives an indication of how efficiently *k*-cliques are put together around *v* to create (*k* + 1)-cliques. This number can be shown to be arbitrarily small or large, depending on the ambient graph (Levi et al., 2020).

Therefore, to obtain a distribution as a number between 0 and 1, we normalise and define the *k-th density coefficient for a vertex v in a graph* 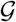 *with n vertices* to be

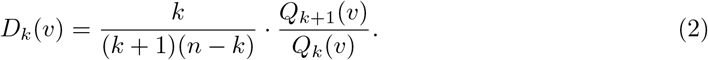

With this definition, if 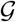 is a directed graph on *n* vertices, where any pair of vertices is reciprocally connected then the *k*-th density coefficient of any vertex in it will be 1 for all *k*. For instance the 2nd density coefficient of a vertex *v* is defined by

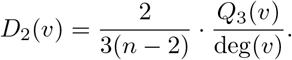

In all cases the subscript *k* refers to (*k* + 1)-cliques being topologically *k*-dimensional. Notice also that *Q*_2_(*v*) = deg(*v*).

### 5.6 Homological parameters

If 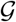 is a directed graph, let 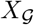 denote its directed flag complex. This is a topological space obtained from 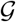 by replacing every directed 2-clique (a directed edge) by a line segment (or in mathematical language - a 1-simplex), every directed 3-clique by a solid triangle (a 2-simplex), every directed 4-clique by a solid tetrahedron (a 3-simplex), and so on in higher dimension. In general each directed (*n* + 1)-clique is replaced by an *n*-simplex - an *n*-dimensional object that generalises the concepts of, line segment, triangle and tetrahedron to a general dimension. The *directed flag complex 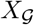 of the directed graph* 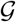 has been defined and studied in the context of neuronal networks and neuroscience in Reimann et al. (2017) and Masulli and Villa (2016).

Any directed (*n* + 1)-clique contains exactly *n* + 1 directed *n*-sub-cliques. The corresponding *n*-simplex contains *n* + 1 (*n* − 1)*-dimensional faces*, that are themselves (*n* − 1)-simplices in 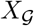. Using this structure it is possible to associate with the directed flag complex an algebraic object called a *chain complex*. It is constructed as follows. For every *n* ≥0, take 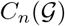 to be the vector space over the finite field of 2 elements (or any other field) Lidl and Niederreiter (1997), with basis given abstractly by the set of all *n*-simplices in 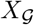 or equivalently all the directed (*n* + 1)- cliques in 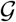. Taking faces of simplices gives rise, for every *n* ≥ 1, to a linear transformation called a differential

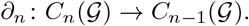

The *n-the homology group* of 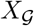 (with coefficients in the field of 2 elements) is defined by

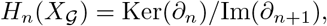

namely the quotient vector space of the kernel of *∂_N_* by the subspace that is the image of *∂_n_*_+1_. For the purpose of the discussion in this article, what is important to understand is only the dimension of the vector space 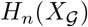 - the so called the *n-th Betti number of* 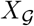. If we let *D_n_* denote the matrix representing the linear transformation *∂_n_*, then the *n*-th Betti number, denoted *b_n_* or 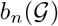, is calculated as the difference

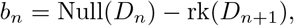

i.e., the dimension of the null-space *D_n_* minus the dimension of the column space of *D_n_*_+1_. These numbers depend on the field as well as on the graph 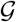, but in this article we worked only with the field of 2 elements. The interested reader is referred to Hatcher (2002) for a thorough mathematical background on these concepts.

The 0-th Betti number *b*_0_ is exactly the number of connected components of 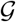. The 1-st Betti number *b*_1_ can be thought of as the number of distinct loops in 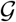 that are not bounding a 2-dimensional subspace in 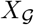. Intuitively, the Betti numbers of 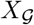 are a count of *n*-dimensional “cavities” in 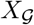 (Reimann et al., 2017).

In this paper we consider two extra topological metrics that are associated to Betti numbers. The first is the classical *Euler characteristic.* The Euler characteristic 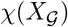 can be computed in two ways that yield the same number. One is the alternating sum

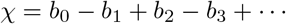

of the Betti numbers of 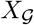. The other is the alternating sum of the number of directed cliques in each dimension in 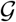. The Euler characteristic is relatively very easy to compute (using the second method) and although it is considered to be a weak invariant of topological spaces, it is frequently extremely efficient both in theory and in applications. It is used in a classification task in Levi et al. (2020) and as a topological parameter in this article.

The second topological parameter we use here is the *normalized Betti coefficient*. It is a weighted sum of the Betti numbers:

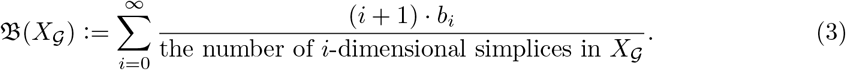

The Betti coefficient is a rough measure of how efficient the graph 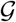 is in creating cavities in all dimensions where they exist in the directed flag complex 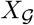.

### 5.7 Spectral parameters

Every square *n × n* real valued matrix *A* has *eigenvalues* {λ_1_*, …, λ_n_*}, which are the (real or complex) solutions to the *characteristic equation A***x** = *λ***x** where **x** is a vector of length *n*. Equivalently the eigenvalues are the roots of the *characteristic polynomial of A*. The collection of eigenvalues of a matrix *A* is often referred to as the *spectrum* of *A*.

Considering the set of moduli (absolute values) of the eigenvalues of *A*, one associates three invariants with *A*:

- The *spectral radius* of *A* is the largest modulus of an eigenvalue.
- The *low spectral gap* of *A* is the smallest modulus of a non-zero eigenvalue of *A*.
- The *high spectral gap* of *A* is the difference between the moduli of its two largest eigenvalues (sorted by their moduli).

Let 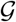 be a directed graph with *n* vertices. The *adjacency matrix A* of 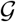 has (*i, j*)-entry *a_i,j_* = 1 if there is a directed edge from vertex *i* to vertex *j*, and 0 otherwise. The *Chung– Laplacian matrix* has a more involved definition, which can be found in Chung (2005). A graph is said to be *strongly connected* if for any two distinct vertices *u* and *v* in 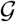 there is a directed path in 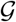 from *u* to *v*. The Chung–Laplacian matrix is only defined on a strongly connected graph. In this paper we consider the spectral radius of the adjacency matrix of the graphs we studied, as well as its low and high spectral gaps. We also considered the Chung–Laplacian spectral gaps of the largest strongly connected components of graphs, whenever these can be uniquely determined.

These invariants are well studied in theoretical and applied graph theory, where their main objective is to relate various structural properties of graphs to the spectrum. For example the Laplacian spectral gap is famously known to measure how easy the graph is to disconnect through so-called Cheeger inequalities (Chung, 1996).

### 5.8 Relative boundary

For a subset *S* ⊆ *V* of vertices in a graph 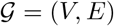, the *edge boundary*

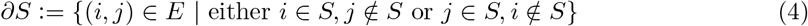

of *S* consists of all edges with exactly one endpoint in *S*. Likewise, the *edge volume*

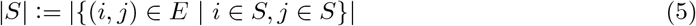

of *S* is the number of edges whose both endpoints are in *S*. We then define the *relative boundary*

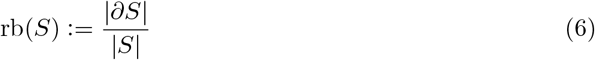

of *S* as the size of its edge boundary divided by its volume. Relative boundary is related to the so-called isoperimetric or Cheeger number (Chung, 1996) and is designed to measure how strongly the subset *S* connects to the rest of the graph relative to its internal connectivity.

### 5.9 Extension

For a subset *S ⊆ V* of vertices in a graph 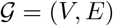, its extension is the number of vertices that are connected to *S* but are not in *S* itself. The *afferent extension* of *S* and the *efferent extension* of *S* are defined as

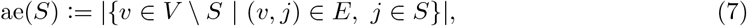

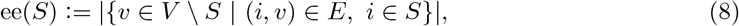

respectively. Note that, for example, a vertex *v* outside of *S* for which (*v, s*_1_), (*v, s*_2_) ∈ *E* for distinct *s*_1_*, s*_2_ ∈ *S* will be counted only once in ae(*S*), so as to distinguish the extension of *S* from the edge boundary of *S*. In other words, ae(*S*) + ee(*S*) is bounded above by the edge boundary of *S*, and the bound is attained whenever no vertex in *V S* is connected by more than one edge to *S*. In that case, the edge boundary coincides with the extension.

### 5.10 Analysis of simulation results

The results of the simulations were analyzed according to the pipeline depicted in Fig. 1. Inputs were:

1. The spike trains of the 31,346 most central excitatory neurons in the simulations
2. For each neuron its morphological type, layer and location in the model (x,y,z-coordinates)
3. The adjacency matrix of synaptic connections between all neurons in the model
4. The identifier of the pattern presented during each stimulation (the “stimulus stream”).

These inputs can be downloaded from https://doi.org/10.5281/zenodo.4290212. The analysis pipeline conceptualized in Fig. S1 was implemented in custom python code (python version 3.7), except for parts of the topological analysis which used custom c++ code that was wrapped for use in python using pybind11. The entirety of the pipeline can be obtained from: https://github.com/BlueBrain/topological_sampling.

The following sections will explain the individual steps of the pipeline.

**Supplementary Figure 1:**
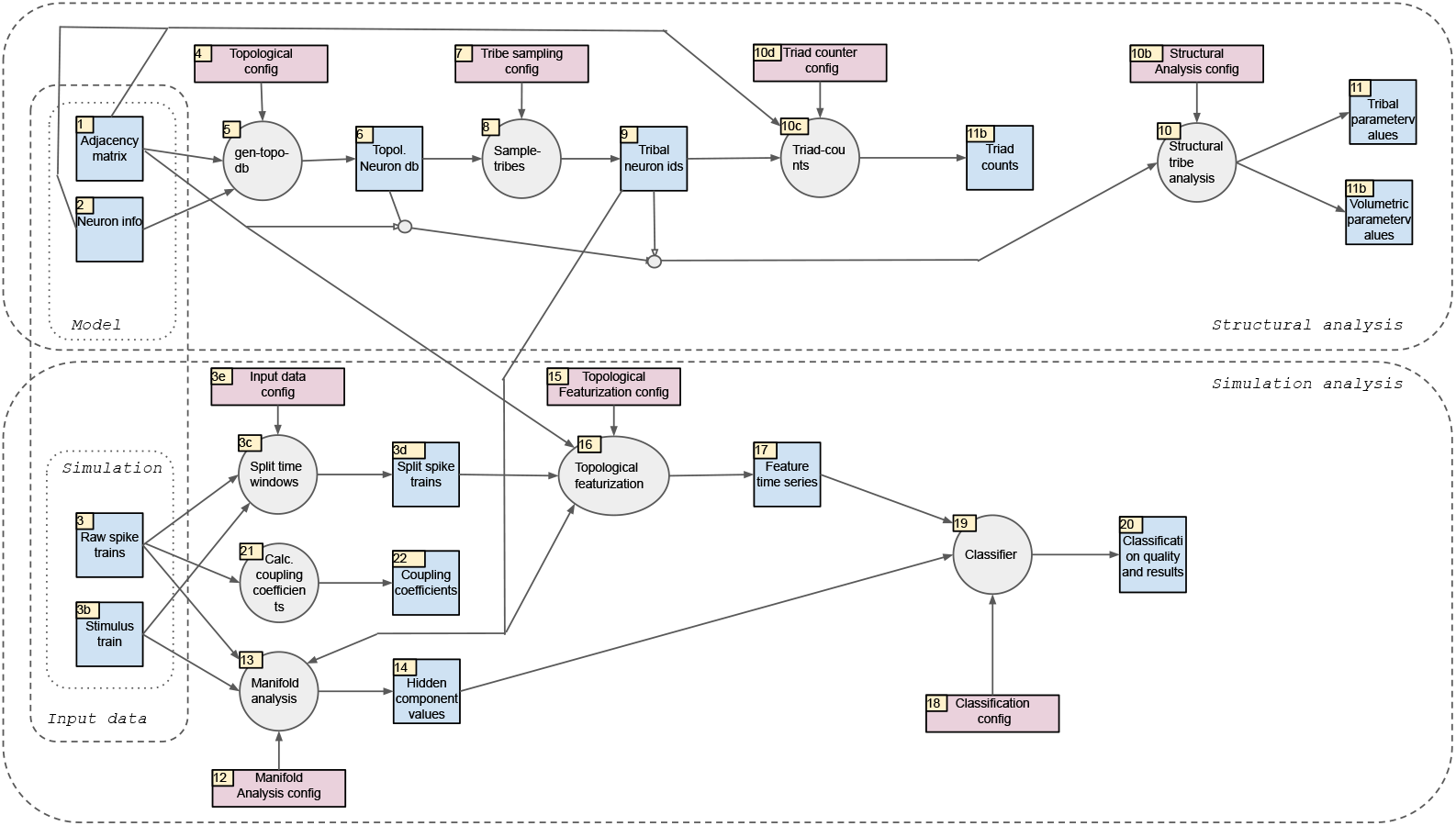
Overview of the inputs into the analysis pipeline and its individual analysis steps.

### 5.11 Generating neuron samples (Sample-tribes)

Neuron samples were generated in one of five ways:

1. Volumetric. First, we found the center of the neuron population by averaging their x, y, and z-coordinates. Next, we added a random offset between 100*μm* and 100*μm* for the x- and z-coordinates (parallel to layer boundaries) and 300*μm* and 300*μm* for the y-coordinate (orthogonal to layer boundaries). Next, we found all neurons in the model within a certain radius of that point and randomly picked 600 neurons from them. We repeated these steps to generate 25 samples each for radii of 125*μm*, 175*μm*, 225*μm*, 275*μm* and 325*μm*.
2. Champions. We generated champions by finding the 25 tribes that yielded the largest values for one of the topological parameters described above. We limited the selection to tribes with at least 50 neurons.
3. Random samples. For each of the morphological types of neurons in the model, we randomly picked 25 neurons and used their associated tribes.
4. Subsampled. For the champions of in-degree, out-degree, and euler characteristic we generated random subsamples. First, we randomly picked five champions of each of the three parameters. Next, we randomly selected a certain fraction of the neurons contained in the tribes. We repeated this five times, generating five subsamples of each picked tribe. We thus generated subsamples at 90%, 70%, 50%, 25% and 15% of the original tribe sizes.
5. Subtribes. For volumetric samples, we picked the 25 largest tribes contained within a sample.

### 5.12 Triad counts

Input into the triad counter was the adjacency matrix of the model and the identifiers of neurons in a given sample. First we extracted the submatrix defining the internal connectivity of the sample. Then we recursively iterated through all possible combinations of three neurons of the sample and classified their connectivity into one of 13 motifs. We limited this analysis to neuron triplets that were strongly connected in either direction, otherwise three additional motifs (unconnected, single connection, single bidirectional connection) would have been possible.

We then calculated the expected numbers of each motif in two control models. First, an Erdos-Renyi graph with the same number of nodes and edges, yielding *C_ER_*(*i*), the expected count of motif *i*. Second, an Erdos-Renyi graph with the same number of nodes and edges, but taking into account the “tribal” sampling procedure. That is, taking into account that the sample contains one central neuron and its neighbourhood. We calculated this as follows.

First, we calculated the number of triplets that contain the chief (central neuron):

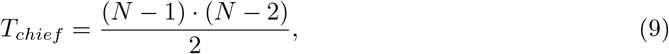

Where *N* refers to the size of the sample. In this arrangement, the connections binding the chief to the two other neurons can have one of three patterns. From the view of the chief they can be efferent-efferent (probability 0.25), efferent-afferent (probability 0.5) or afferent-afferent (0.25). For each of these possibilities we analytically derived the expected numbers of motifs assuming that the remaining connections were subject to a uniform, statistically independent probability derived. This included the option to turn the connection binding the chief to a tribe member into a bidirectional connection. The thusly derived motif probabilities, *P_chief_*, multiplied by the number of chief-including triplets yielded the first part of the expected motif counts.

Next, the number of triplets not containing the chief was:

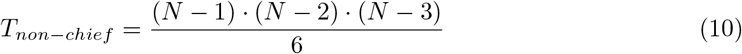

For these we derived the expected motif counts according to an Erdos-Renyi control. Total expected count of motif *i* in tribal sampling was then:

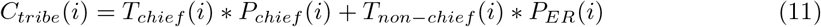

The degree of over- and under-expression of motif *i* in volumetric samples was then calculated as:

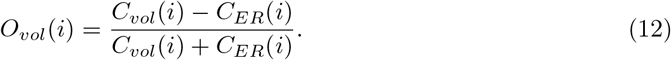

The degree of over- and under-expression in champion samples was calculated relative to the volumetric samples. First we normalized each sample against their respective control:

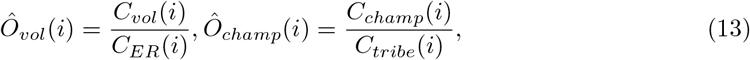

Then, we normalized the motif count in the samples to the mean and standard deviation of the volumetric samples:

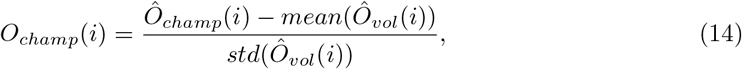

Where *mean* and *std* refer to the mean and standard deviation over volumetric samples.

### 5.13 Calculating topological parameters for samples

We calculated the values of the topological parameters for all possible tribes, i.e. we considered each neuron in the model as a chief and calculated the parameters of the resulting tribe. We also calculated parameter values for volumetric samples in two ways.

First, by simply applying the topological method to the connectivity of the sample. This could be done for all parameters except in-degree, out-degree, transitive clustering coefficient. These parameters were calculated in tribal samples as the in-degree, out-degree, etc. *of the chief* and were consequently undefined for volumetric samples. We instead used the mean in-degree, out-degree of clustering coefficient of all neurons in the volumetric sample.

Second, by taking the tribal structure of the sample into account. We did this by calculating the relative overlap of the sample with each possible tribe in terms of neuron contents. Then we calculated the value of a topological parameter as a weighted average of the *n* strongest overlapping tribes with the weight being proportional the size of the overlap. We optimized the value of *n* to yield the best predictor of accuracy, separately for each topological parameter.

### 5.14 Calculation of coupling coefficients

Input into this step were the raw spike trains of all neurons recorded in the simulations.

We started by binning the spike trains of all neurons in the model into 10 ms bins. This yielded a *N × T* sparse matrix, where *N* was the number of neurons and *T* the number of time bins and the entry at *i, j* specified the number of spikes of neuron *i* in time bin *j*. For the coupling coefficient of neuron *i*, we used the *i*th row of the matrix as the time series of firing of that neuron, and the mean value of all rows of the matrix, except *i* as the average firing rate of all others. We then calculated the coupling coefficient as the normalized correlation between the two time series (numpy.corrcoef).

### 5.15 Manifold analysis

Input into this step were the raw spike trains of all neurons recorded in the simulations, information about the pattern identity of each stimulation, and the identifiers of a sample of neurons. We began by binning the spike trains of all samples neurons into 10 ms bins. This yielded a *N_sample_ × T* sparse matrix, where *N_sample_* denotes the number of neurons in the sample. Non-spiking neurons were removed. Next, we extracted the twelve strongest components from this *N_sample_*-dimensional time series using factor analysis (sklearn.decomposition.FactorAnalysis). Then we split the resulting 12-dimensional time series into 200 ms time windows that each corresponded to a stimulus presentation. Finally, we grouped these time windows by the identity of the stimulus pattern presented at that time.

### 5.16 Stimulus classification

Input into the classifier was either: The twelve strongest components of the spiking activity of a tribe, extracted as detailed above. Or: The time series of euler characteristic values of the active sub-tribe extracted from the activity of 25 tribes as detailed in the main text. In both cases, the input was split into time windows that each represented a single trial, i.e. a single stimulus. The time windows were further grouped by the identity of the stimulus pattern used in the trial.

Next, we generated for each time window a time series of expected outputs of the classifier. This was simply the identity of the stimulus (an integer between 0 and 7), repeated for each time step. 60% of the trials and their associated expected outputs were used to train a linear classifier (sklearn.svm.SVC). The remaining 40% were used to determine classification accuracy. This was repeated 5 times, thereby conducting 6-times cross validation.

### 5.17 Dependence of classification accuracy on topological parameters

We first generated a model of accuracy on tribe size. To that end, we calculated for each sample its *mean tribe size*, i.e. the mean over all neurons in a sample of the size of the tribe associated with the neuron. This could be calculated equally for tribal and volumetric samples. Next, as the model of the impact of tribe size, we conducted a linear fit of *mean tribe size* against classification accuracy (using the manifold method) based on the data of 1276 randomly picked tribes. The fit minimized the sum of squared errors and was performed using the statsmodels package in python 3.7. Specifically, we used the formula based interface with the following formula:

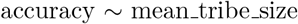

We calculated the *residual accuracy* for each sample – tribal or volumetric – by subtracting the prediction of this model from the classification accuracy values.

We then determined the relation between topological parameters and residual accuracy as follows. We began with a simple control model predicting the residual accuracy from the sampling radius in the case of volumetric samples or from the morphological type of the chief in the case of tribal samples. Both morphological type and radius were considered categorical variables. We obtained the models with the formula based interface to statsmodels as:

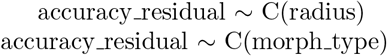

Next, we normalized the values of each topological parameter to zero mean and unity variance and generated linear fits taking into account the effects of both radius / morphological type and and individual, normalized parameter value:

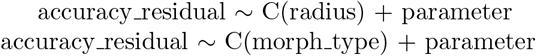

We then calculated the slope of the fits in the parameter to assess the strength of the effect of that parameter; and the fraction of variance explained, from which we subtracted the fraction of variance explained by the simpler control models.

## 6 Author contributions

Conceptualization, M.W.R., R.L., H.R.; Methodology, M.W.R., H.R., J.S.; Software, M.W.R., H.R., J.S. J.L., C.P.; Validation, M.W.R.; Investigation, M.W.R., J.S., H.R., C.P.; Visualization, M.W.R., J.L., C.P.; Writing – Original Draft, M.W.R.; Writing – Review & Editing, M.W.R., R.L., J.L., H.R., C.P.; Supervision, M.W.R., R.L.

## 7 Acknowledgements

This study was supported by funding to the Blue Brain Project, a research center of the École polytechnique fédérale de Lausanne (EPFL), from the Swiss government’s ETH Board of the Swiss Federal Institutes of Technology.

Levi is supported by an EPSRC grant EP/P025072/ and a collaboration agreement with École Polytechnique Fédérale de Lausanne.

## 8 Data and code availability

The data used is available at https://doi.org/10.5281/zenodo.4290212.

The entire analysis code can be obtained from: https://github.com/BlueBrain/topological_sampling.

## 9 Supplementary Material

### 9.1 Supplementary explanations

Here we expand on the topological methods presented in Section 5.2 and subsequent sections, and describe the parameters from Table 1 in a more heuristic manner trying to motivate them from a neuroscientific point of view. Figure 2 is a visualization of some of the parameters.

#### 9.1.1 In-degree and out-degree

In signal-flow networks, a high in-degree indicates that a vertex has a potential to receive inputs from many other vertices, while high out-degree indicates potentiality to transmit outputs to many vertices. In a neuroscience context, high in-degree of a vertex *v* indicates that the corresponding neuron is potentially receiving spikes from many neurons, while a high out-degree means the neuron is potentially transmitting its spikes to a large number of synaptically connected neighbours.

#### 9.1.2 Clustering coefficient

The clustering coefficient at a vertex *v* is a measure of how interconnected the vertices of the neighbourhood of *v* are. A “possible” directed 3-clique at *v* is formed by two directed edges containing *v* and two other distinct vertices. This coefficient aims to measure how far away the neighbourhood of *v* is from complete communication, in a directed way, within all its triplets of neighbours.

#### 9.1.3 Density coefficient

Slightly coarser than the clustering coefficient is the 2nd (unnormalized) density coefficient of *v*, which is the number of directed 3-cliques that contain *v*, divided by the degree of *v*. This value is normalized so that *v* in a complete directed graph on *n* vertices has density coefficient 1. The idea is that the larger the density coefficient of *v* is, the more the vertex participates in “informational exchange”, as indicated by the presence of many directed 3-cliques.

**Supplementary Figure 2:**
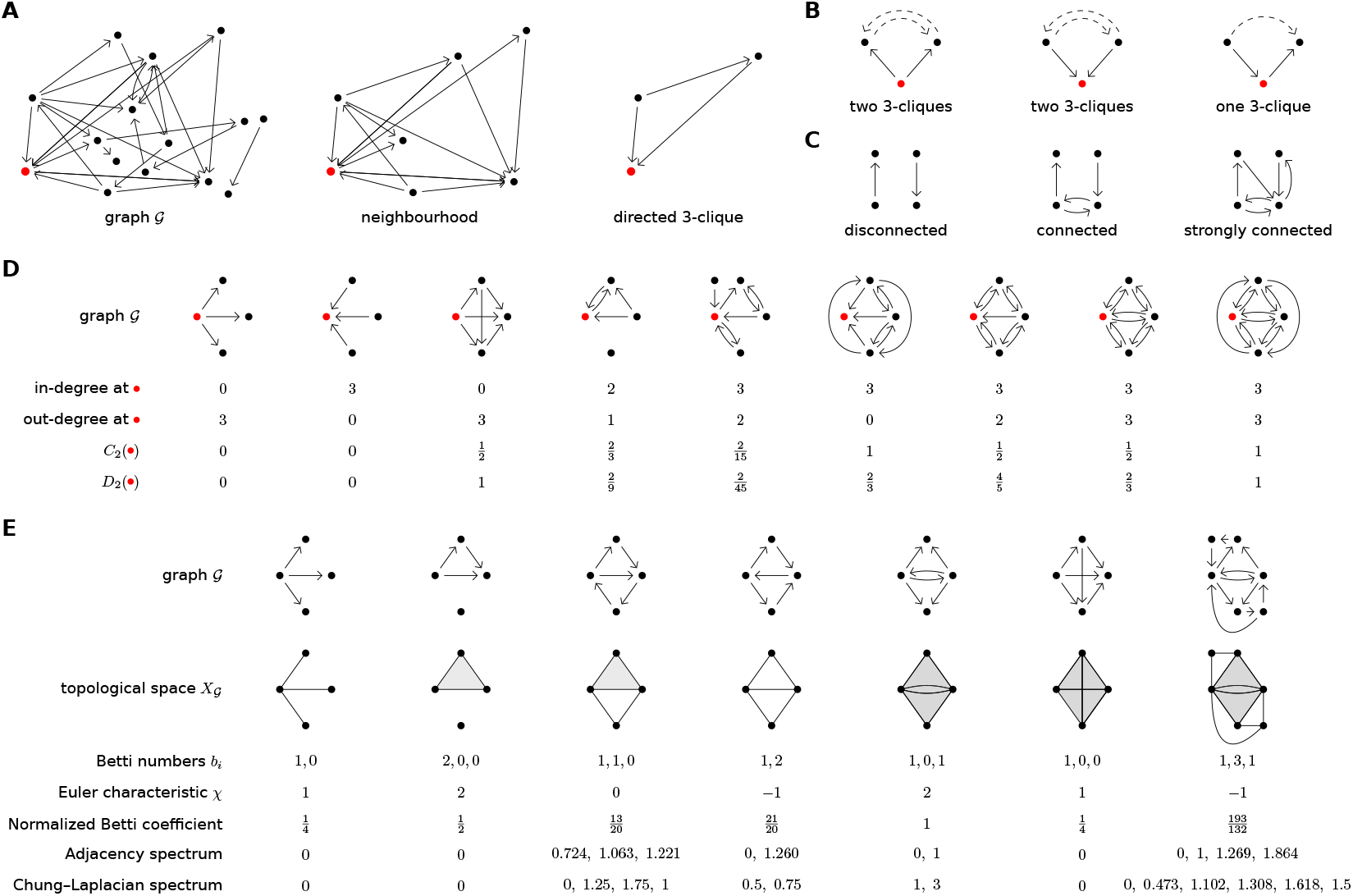
(A) Graphs, neighbourhoods, and cliques. (B) Different ways to complete two edges to a directed 3-clique. (C) Different types of graphs. Note the Chung–Laplacian spectrum considers only the largest strongly connected component when computing eigenvalues. (D) Comparisons of four different graph parameters relative to one of its vertices. (E) Comparisons of three different graph parameters and two different spectra (unique absolute values of eigenvalues of matrices).

#### 9.1.4 Homological parameters

The *Betti coefficient* as compared to the Euler characteristic (Section 5.6) is a more refined method to summarize the Betti numbers, by adding scaled Betti numbers together. This number is normalized, so that it has value 1 if the topological space *X_G_* associated to the graph *G* is a sphere (of any dimension). In other words, as the *n* + 1 Betti numbers for an *n*-dimensional sphere are 1, 0, 0*, …*, 0, 0, 1, the normalized Betti coefficient measures how close *X_G_* is to being a sphere, where “close” is used in terms of Betti numbers, that is, in terms of different dimensional holes.

#### 9.1.5 Spectral parameters

Solution vectors **x** to the characteristic equation *A***x** = *λ***x** are the eigenvectors of the matrix *A.* These eigenvectors in the graph setting are vertex functions. When *A* is the adjacency matrix of a graph, the *k*th element in the product vector *A***x** is the sum of the function values at the vertices that the *k*th vertex has a directed edge to. If this equals the *k*th element of **x** scaled with *λ_i_* as in the characteristic equation, we see that heuristically the eigenvalues of an adjacency matrix can be seen as scalings of vertex values in a “balanced” signal transmission in a sense that a vertex fully distributes its value to its out-neighbours. The spectral radius and spectral gap (low) then correspond to the maximal and minimal transmission and spectral gap (high) to the difference between the two most maximal transmissions. Similar derivation for Laplacian is not so easy but the Laplacian spectral gap is a measure of robustness of the graph in the sense that how difficult the graph is to disconnect into two separate sets of vertices by removing edges 5.7.

**Supplementary Figure 3:**
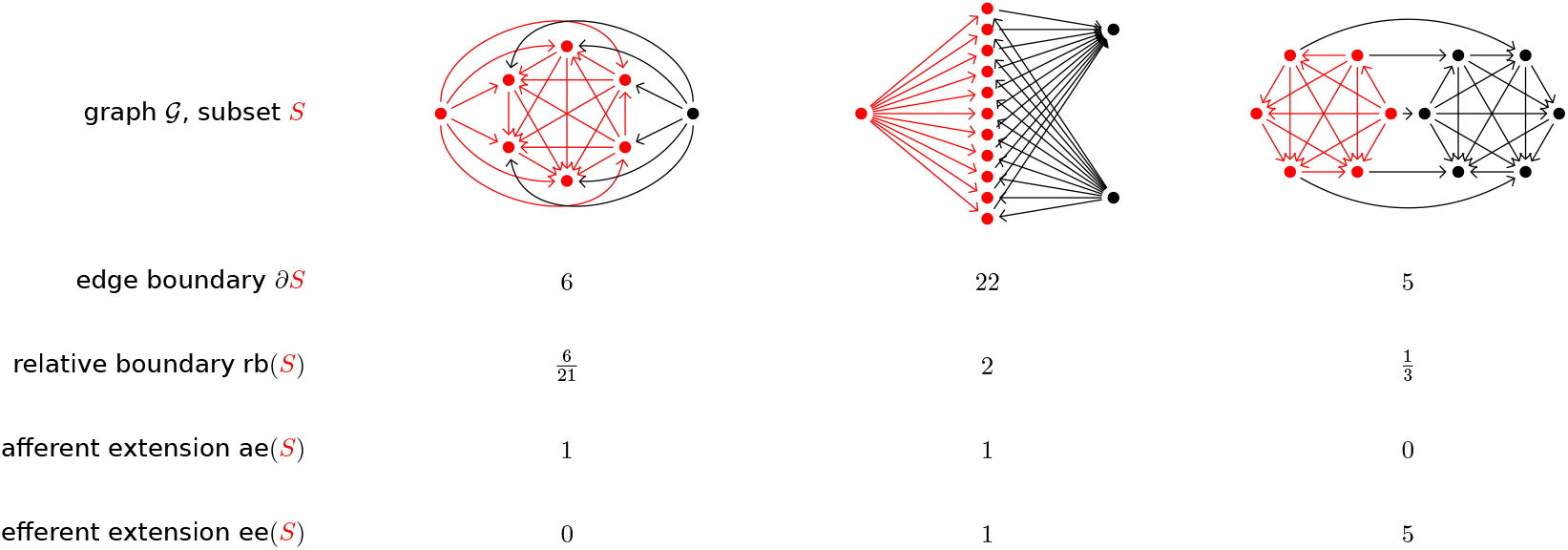
Comparisons of the edge boundary, relative boundary, afferent extension, and efferent extension for particular subsets. Subsets are chosen as tribes.

#### 9.1.6 Relative boundary

A subset of vertices in a graph might connect within themselves while also connecting to vertices not in the subset. Relative boundary therefore measures the relative dominance between these two connectivities. It is a measure of the communication potential within a subset, in the same fashion as the clustering coefficient, while also taking into account communication with the ambient graph.

#### 9.1.7 Extension

The afferent extension is defined to measure how many different vertices in the ambient graph might send information into the selected subset of the graph. Likewise the efferent extension measures how much information the subset can potentially send to the ambient graph. Neural spikes flow between neurons through directed synaptic connections and the amount of spikes that can pass through a neuron depends on how many neurons it is connected to. The extension aims to mimic this for a collection of neurons, separating it to incoming and outgoing flow.

### 9.2 Supplementary figures

**Supplementary Figure 4:**
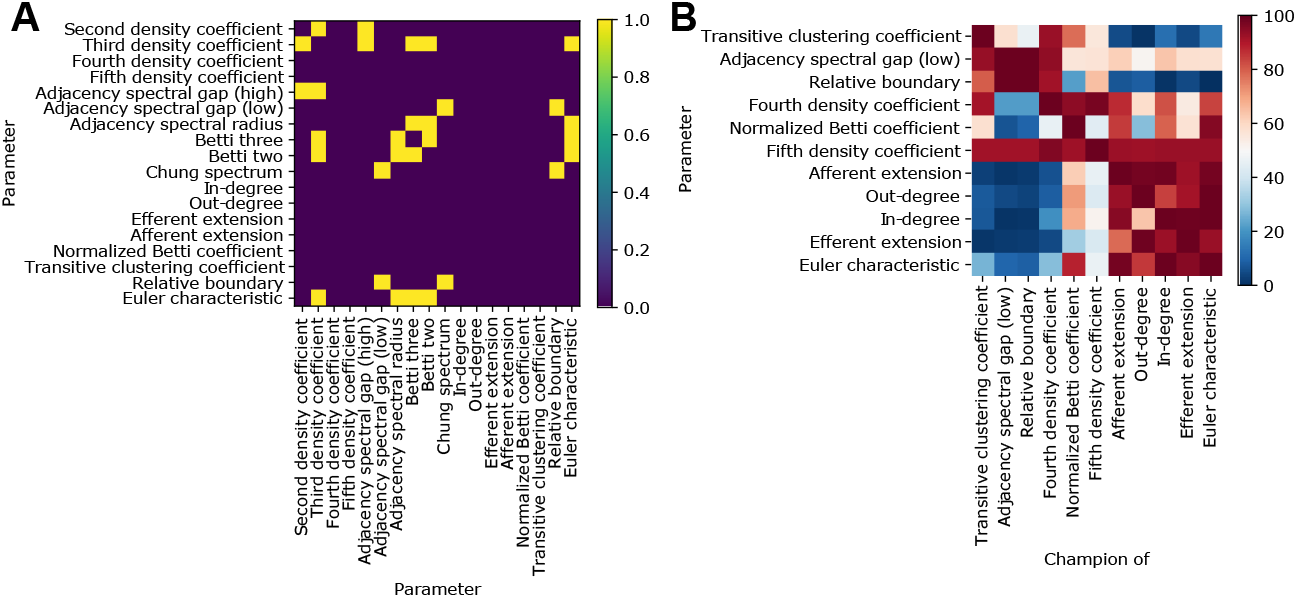
A: All investigated topological parameters, with pairs that are mutually redundant (in terms of resulting triad motif expression patterns) highlighted in yellow. B: For the champion tribes of the non-redundant parameters (columns), we consider the values of all parameters (rows), normalized in terms of the percentile of the overall distribution of said parameter (see color bar).

**Supplementary Figure 5:**
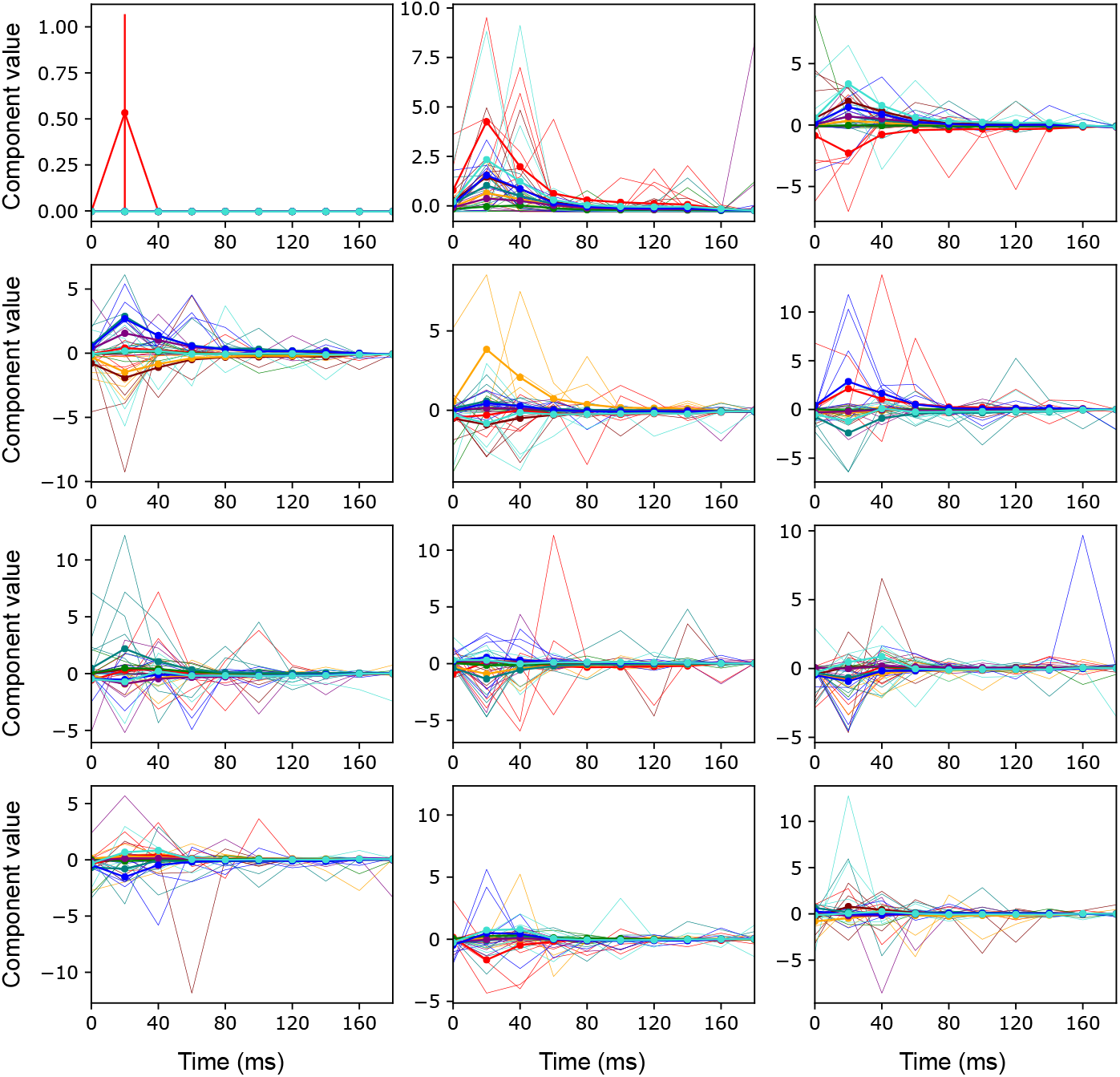
Time series of the twelve strongest components (panels) during presentation of the individual stimulus patterns (colored traces). Thick lines and error bars: mean and SEM. Thin lines: for five randomly selected trials using a given pattern.

**Supplementary Figure 6:**
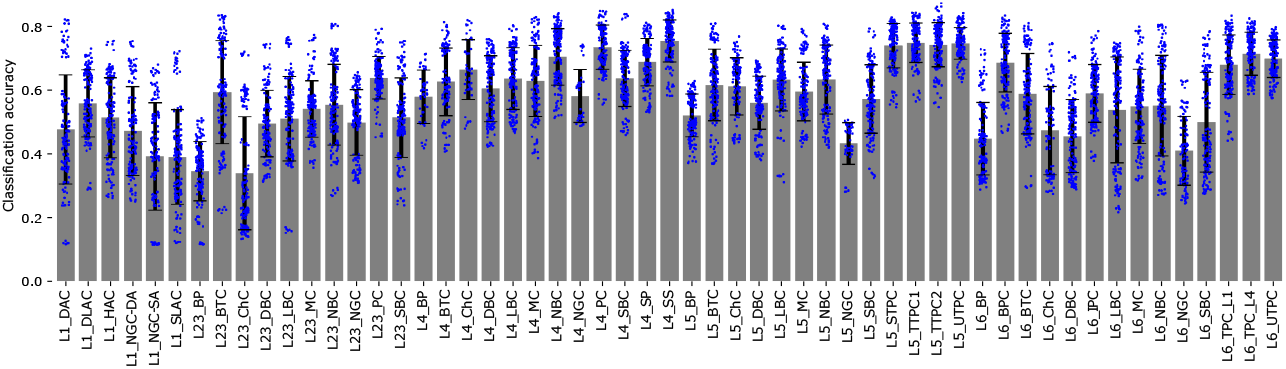
Classification accuracies, using the manifold-based method, for the randomly selected tribes with chiefs of the indicated morphological type. Grey bars and error bars: mean and std. Blue dots: individual tribes.

**Supplementary Figure 7:**
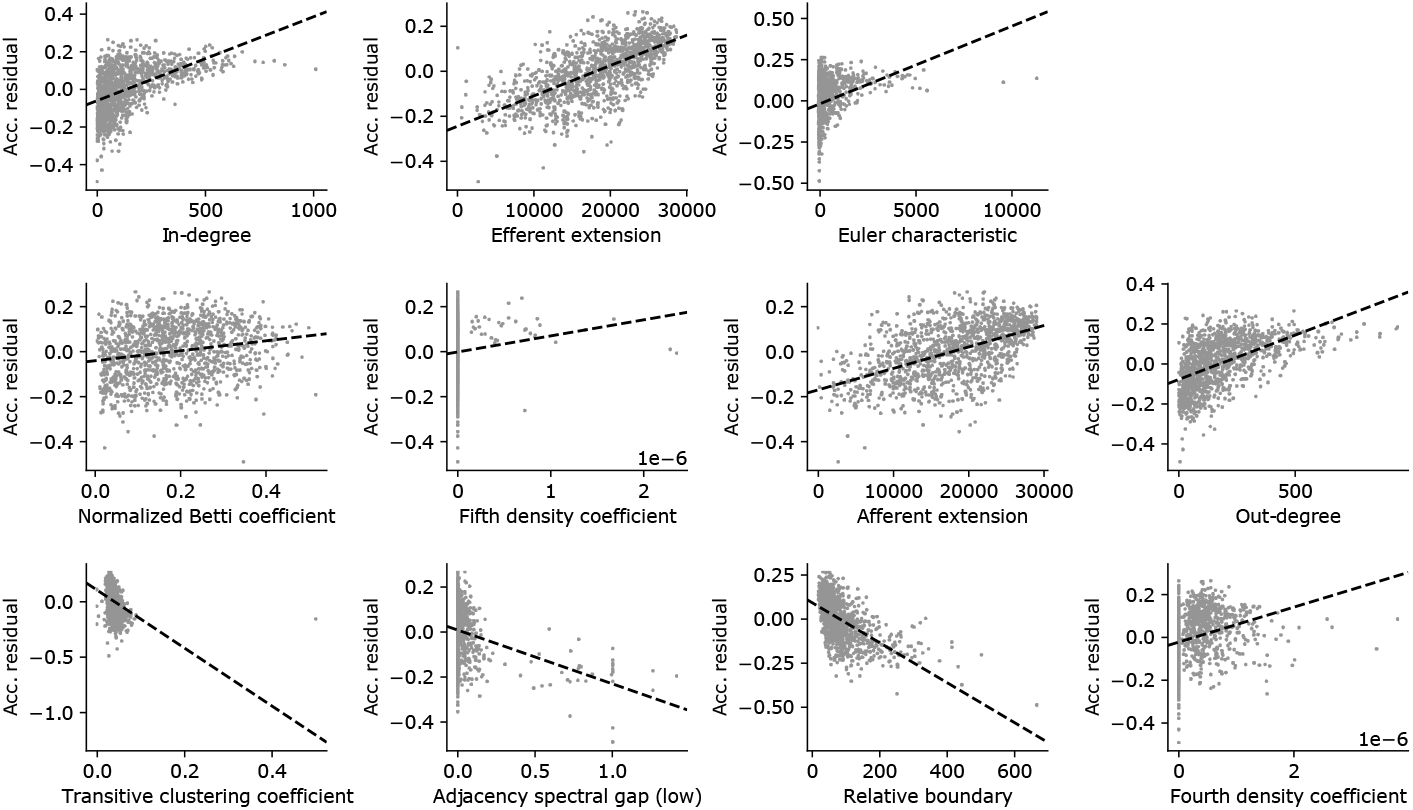
Values of topological parameters against residual accuracy for randomly selected tribes. Grey dots: individual tribes. Black line: linear fit.

**Supplementary Figure 8:**
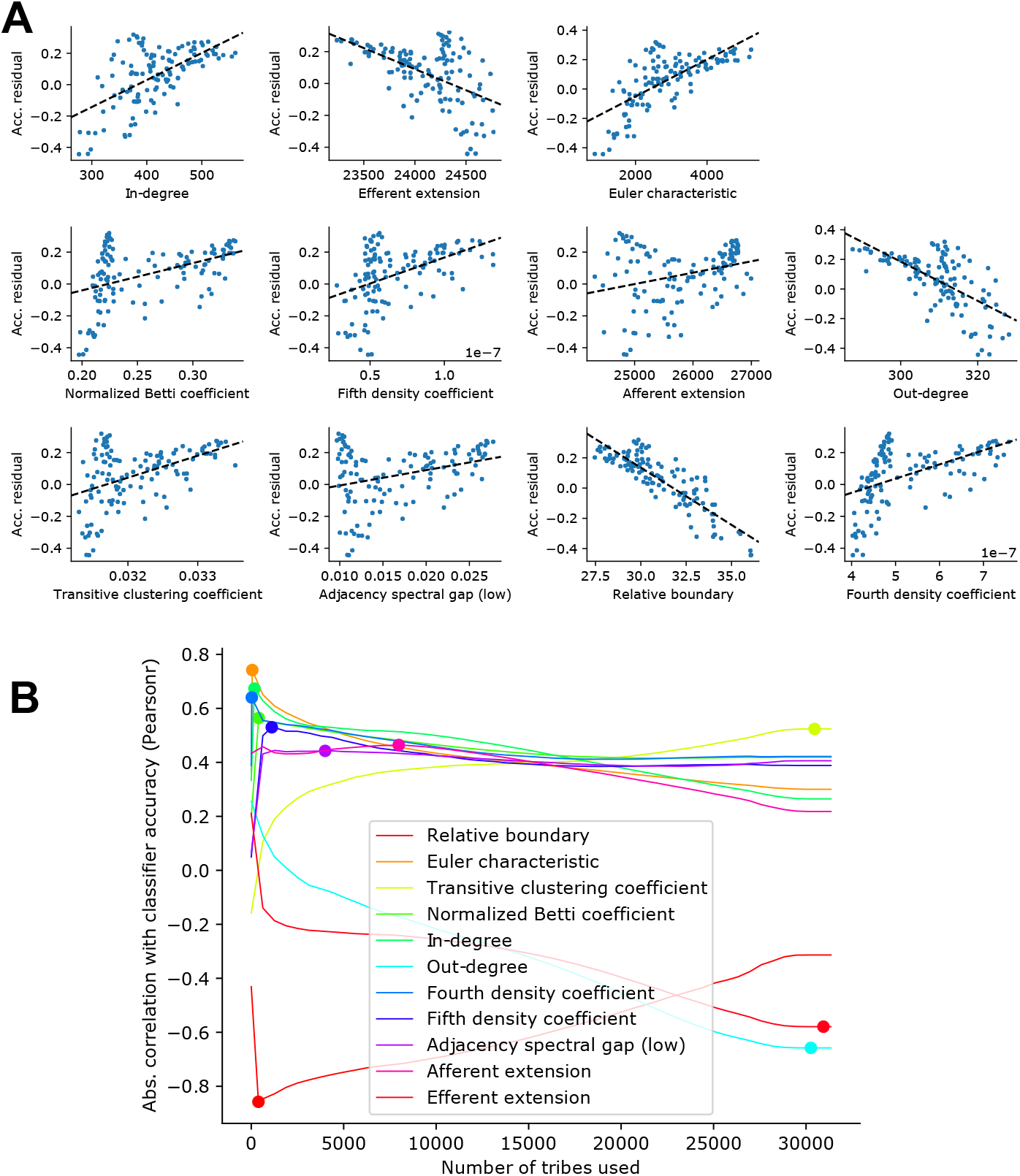
A: Synthetic values of topological parameters for volumetric samples against their residual accuracy. Blue dots: individual samples. Black line: linear fit. B: Number of tribes used in the calculation of the synthetic values (see Sec. 5.13) against the resulting correlation (pearsonr) with classifier accuracy. Individual, colored lines: For individual topological parameters. Colored dots: Maxima of the absolute value of correlations, indicating the number of tribes used in the remainder of the manuscript.

**Supplementary Figure 9:**
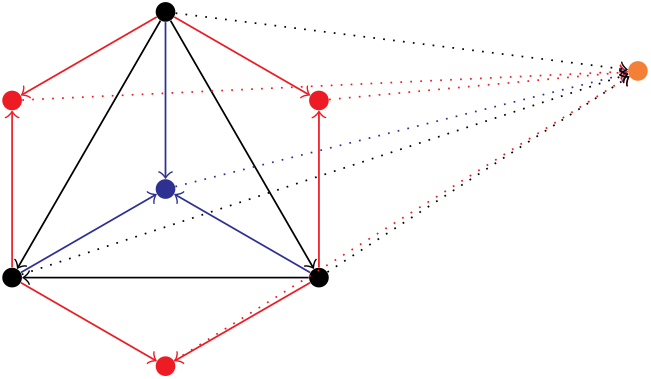
The neural circuit to read out the Euler characteristic of the active circuit of a directed 3-clique, the graph in solid black. The red neurons are the read out neurons for the 2-cliques, the blue neuron is the read out clique for the 3-clique. The black and blue dotted lines have weight 1, and the red dotted lines have weight 1. The orange neuron stores the Euler characteristic of the active subgraph.

